# Nonspecific interactions can lead to liquid-liquid phase separation in coiled-coil proteins models

**DOI:** 10.1101/2025.05.09.653163

**Authors:** Dominique A. Ramirez, Anastasia Shrimpton, Michael R. Shirts, Loren E. Hough

## Abstract

Liquid-liquid phase separation (LLPS) is one mechanism that cells can use to organize biomolecules spatially and functionally. Some coiled-coil (CC) proteins, such as the centrosomal proteins pericentrin and spd-5, are thought to LLPS, but it is currently unknown what parts of these proteins facilitate the process. It is thought, however, that the numerous CC domains in these proteins might be contributing to their LLPS. We recently showed, using computational studies and designed proteins, that CC domains can facilitate LLPS through specific interactions between the CC domains themselves, meaning that each CC was designed to interact only with a subset of other CCs in the system. This is in contrast to nonspecific interactions, where all CCs would be able to interact with all other CCs in the system, which is akin to some interactions (e.g. *π*–*π*) seen in phase-separating intrinsically disordered proteins. Because the specificity of interactions between natural CC domains is tunable in a sequence-dependent fashion, CC domains present a unique system that allows us to investigate the contributions of specific versus nonspecific interactions on LLPS. We show, in our computational system, that CC proteins with nonspecific interactions can LLPS but with less propensity compared to specific interactions. The LLPS propensity of CC proteins with nonspecific interactions can be improved by altering the structure and dynamics of linker segments, without directly changing the specificity of interactions. We also demonstrate that the number of intra-chain CC contacts plays a direct role in determining LLPS for nonspecifically interacting proteins. These results have broad implications for the role of linker segments—protein features beyond the interaction domains e.g. ‘stickers’—in protein LLPS and the formation of biomolecular condensates.

**STATEMENT OF SIGNIFICANCE:** Model coiled-coil proteins, which use coiled-coil domains as stickers, are capable of phase separation in a regime where intra-protein contacts interfere with the interactions which support phase separation. We explore ways to increase phase separation propensity without changing interaction specificity and find that the structure and size of spacers impacts LLPS propensity by affecting the formation of intra-chain interactions. This work demonstrates that protein LLPS might be controllable without directly affecting the cohesive parts of a protein i.e. stickers. This work also suggests that LLPS propensity might be a broadly accessible phenomenon for coiled-coil proteins.

## INTRODUCTION

The liquid-liquid phase separation (LLPS) of biological polymers plays an important role in cellular physiology. LLPS is a process in which a dissolved polymer can, at a critical concentration, demix from solution to form a polymer-rich liquid phase that coexists in a polymer-diminished solvent. Many proteins and other biological polymers such as mRNA have the ability to LLPS, and when they do so in a cellular environment, they can form biomolecular condensates. Biomolecular condensates can serve many cellular purposes, including sequestering specific molecules out of the cytosol (1), or acting as reaction centers for enzymes (reviewed further by LaFlamme and Mekhail (2), and O’Flynn and Mittag (3)). LLPS is also thought to be an important mechanism for the formation of functional membraneless organelles (4), including stress granules, P-bodies, nucleoli, and centrosomes (5). There is a putative link between altered LLPS propensity of some proteins and certain diseases such as neurodegeneration and cancers (6–8), but the pathological implications of aberrant LLPS are not well characterized. A better understanding of biological LLPS, and the conditions under which it is physiologically relevant, might aid in the treatment of human diseases associated with misregulated LLPS.

Proteins that LLPS can be thought of like associative polymers—they contain *stickers* which are the driving motifs for LLPS, and *spacers* (or linkers) which act to separate stickers within a single protein chain but can still impact LLPS propensity (9–11). Protein stickers can be individual amino acids (akin to monomer stickers in the original theoretical framework of associative polymers (9)) but, importantly, stickers can also be structured protein domains (examples reviewed by Mohanty et al. (12) and Tibble et al. (13)) or even patches along a folded globular surface (11, 14). Proteins that LLPS will contain multiple stickers, regardless of the type, that enable them to make multivalent interactions as multivalency is a general feature of any phase-separating protein (14–16). The flexibility of protein spacers can also contribute to a protein’s ability to LLPS, and it is expected that some amount of disorder is important for LLPS, especially when protein stickers are structured domains (17). Indeed, the majority of identified proteins exhibiting LLPS are either intrinsically disordered proteins (IDPs) or have disordered regions (IDRs) that enable conformational flexibility (18–20) and as a result the field has largely concentrated its efforts to study disordered proteins.

Coiled-coil (CC) domains are one example of a structured domain that is thought to enable protein LLPS. CC domains are protein motifs that can interact with each other in a sequence-dependent manner (21–23) to form alpha-helical multimers in which individual domains coil around each other (21). CC proteins can contain one or multiple of these CC domains—for example, proteins in the centrosome family often contain multiple CC domains (24, 25). Simulations of model CC proteins and *in vivo* experiments with engineered CC proteins show that CC domains can drive protein LLPS (26–31), and this is because both CC domains and CC proteins can form multivalent interactions necessary for LLPS. The role of CC domains in the LLPS of natural proteins, however, not yet well understood. Naturally occurring CC proteins across various species have recently been identified as potentially phase-separating, with some examples including pericentrin (32), spd-5 (33, 34), centrosomin (35), GM130 (36), and EDE1 (37). A few studies have implicated CC domains in the LLPS of natural CC proteins (34, 37–39), but work remains to provide more direct evidence.

We have developed a simulation framework (28), motivated by recently identified phase-separating CC proteins, to study CC-driven LLPS, whereby we treat helical CC domains as stickers and the unstructured regions connecting CC domains together as spacers. Our initial simulation study (28) showed that multimeric multivalency (the size of CC multimers, i.e. dimers, trimers, tetramers) has a stronger positive effect on LLPS propensity than polymeric multivalency (the number of coils per protein). We also demonstrated that the helical structure of CC domains was essential for LLPS in our model, suggesting that attractive residue-residue interactions alone are not sufficient to support LLPS and instead need to act cooperatively when appropriately spatially arranged to make CC domains into stickers. We noted that the specificity of sticker interactions (between CC domains) is an unexplored variable in our understanding of CC-driven LLPS, but also more broadly in the context of protein LLPS and warrants further study.

In order to determine the role of specificity in our framework, we define specific interactions when there exists a difference in attractive energy between different sticker types in the system, such that some stickers attract more strongly to certain sticker types than others, depicted schematically in Figure 1. Our classification of specificity is distinct from discussions that focus on the relative strength of protein:protein interactions (40), which are not necessarily correlated with specificity (41). Our definition is also more precise compared to discussions of specificity that focus on the number of different partners a sticker can have, i.e. promiscuity (42), which is instead better described as multivalency. The maximally specific interaction regime, which was the focus of our previous work (28), is such that only inter-chain interactions are allowed (Fig. 1A), and is a regime that readily LLPS’s. Examples of specific sticker interactions include the proteins developed by Li et al. (15) which have specific interactions between PRM and SH3 domains, and engineered coiled-coil domains which interact specifically only with their designed partners (26, 27).

**Figure 1.**
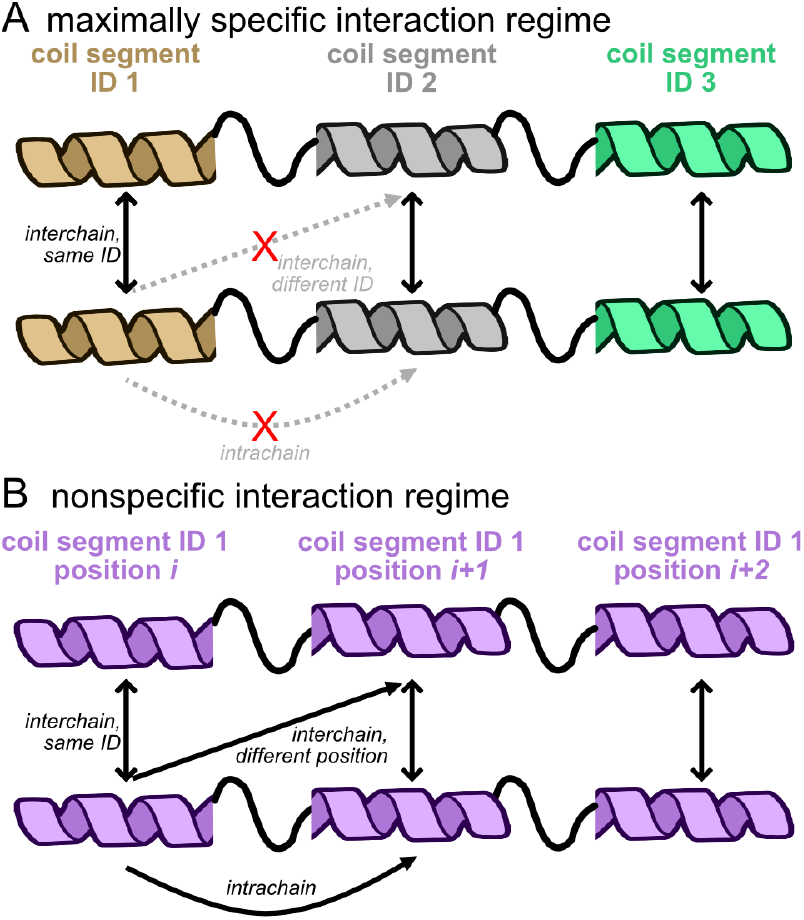
Schematics of protein:protein interaction specificity used in our models of coiled-coil proteins. (A) The maximally specific interaction regime, which was addressed in our previous work (28), is such that each coil segment only interacts favorably with other coils of the same ID (indicated by color in this picture) leading to only inter-chain contacts. (B) The nonspecific regime, where coil segments can interact with equal attractive energy with all other coils, results in inter- and intra-chain contacts. In each diagram, the colored helices represent coil segments and the thick black squiggly lines connecting coils represents the linker segments. Illustration modified from NIAID NIH BIOART Source (bioart.niaid.nih.gov/bioart/468).

Nonspecific interactions, in contrast, are such that stickers have relatively equal interaction strength with all other stickers in the system (Fig. 1B), including those within the same protein chain. It is thought that nonspecific interactions between phase-separating biomolecules could be important for determining LLPS (12, 19, 43–45), but a clear role for nonspecific interactions, if any, is still up for debate (14, 46, 47) Our definition of nonspecificity assumes nothing about the numbers of partners a sticker can have and instead focuses on the energetics of sticker:sticker interactions. Both the specific and nonspecific interaction regimes as described are convenient and simple models to examine, but are ultimately just two extreme limits of protein:protein interaction specificity. Examples of nonspecific interactions and systems can include Coulombic (of opposite charges), *π*-*π* and cation-*π*, interactions between hydrophobic patches, the associative polymers discussed by Semenov and Rubinstein (9, 10), Lennard-Jones toy models used to study phase separation (48), some naturally occurring IDPs known to phase-separate (49), as well complex coacervation between proteins and RNA (50, 51) and DNA (52). An important consequence of nonspecific interactions is that both inter- and intra-chain sticker contacts are possible, because they are enthalpically equivalent, which will change the effective availability of stickers to form inter-chain contacts.

It is unknown how intra-chain contacts between structured domain stickers can affect the LLPS propensities of natural proteins. A study in 2018 showed by simulation that single molecule properties of IDPs—whose stickers are individual amino acids—correlate well with phase separation behavior (53). A conclusion from that study is that in the limit of weak sticker interactions, intra-chain contacts that exist in dilute conditions can rearrange to form LLPS-supporting inter-chain contacts in concentrated solutions (53). Interactions between stickers that are whole protein domains, however, are not expected to be weak and in fact may be high affinity (12), such as between coiled-coil domains (54). It thus becomes important to understand how intra-chain interactions affect LLPS propensity for sticker interactions that might be stronger than those in IDPs, and how these might result in configurations that make inter-chain contacts inaccessible.

We expect that specificity in real proteins is not a binary like the regimes discussed here, but instead is a gradient, with interaction strengths dependent on the amino acid sequences involved. Natural CC domains, for example, are not always expected to interact with high specificity for their cognate partners and can sometimes display nonspecific-like interactions with many similar CC partners (55–57). There could also be nonspecific interactions between CC domains and spacer regions or other structured domains. Additionally, spacer regions may have some relatively weak levels of attraction, but still engage in specific interactions, rather than being entirely unlike the stickers. Because of this gradation across degrees of specificity in realistic proteins, our study of the nonspecific regime might help us understand the ways in which structured domains can enable phase separation of proteins in cases where intra-chain interactions compete for the LLPS-supporting inter-chain contacts.

## METHODS

### Software

All molecular dynamics (MD) simulations were performed using GROMACS (58) v2022.1. Non-parallel-processing GROMACS was compiled on a Ubuntu machine (22.04.2 LTS) with GCC v11.2.0, and on a MacOS 14.6.1 (Intel processor) with AppleClang v.15.0.0 for small-scale simulations and data processing. We compiled a parallel-processing version of GROMACS using GCC v11.2.0 and Open MPI v4.1.1 for use on the Alpine high-computing resource at the University of Colorado Boulder. Custom analysis scripts (available on GitHub, repositories given below) were written for Python 3.9.

### Code availability

Files from this study, including MD parameter files for single molecule generation and slab simulations, relevant topology files (GROMACS .top and .itp), analysis, etc. are available on the GitHub repository associated with this manuscript: https://0020//github.com/dora1300/coiledcoil_nonspecific_llps.git The scripts used to generate proteins following the PeptideBuilder strategy from our previous publication (28) are found at a separate GitHub repository: https://github.com/dora1300/cc_llps_framework.

### Description of the CC-LLPS simulation framework

We previously developed a C-*α* (i.e. one-bead-per-residue) coarse-grained (CG) simulation framework for representing coiled-coil proteins and to study CC-domain-driven LLPS. We use that same simulation framework in the present study. Full design details of the framework are presented in our previous publication (28), but we include a brief description here.

We construct CC proteins as linear combinations of coil segments (representing CC domains) and linker segments, which serve as the stickers and spacers, respectively. This organization allows us to treat CC proteins as associative polymers. Amino acids are represented by a single CG bead centered at the same coordinates as the C-*α* carbon in a corresponding atomistic representation of a protein. Importantly, we treat an entire coil segment as an individual sticker unit, which is made up of CG beads that have their own attractive interaction energy and cooperatively work to create the coil-segment-as-sticker (further description of the interaction parameters provided in Supporting Material sec. S.II). Therefore, the only protein-protein interactions that occur happen between coil segments. Further details about the structural parameters of the framework are included in Supporting Material section S.I.

The focus of the current publication is the *nonspecific interaction regime* (Fig. 1B) in which all coil segments within a single protein and within a simulation box have equal attractive interaction energy. This results in an interaction regime where both inter- and intra-chain coil interactions can occur.

### MD parameters

We used a mostly consistent set of MD parameters throughout the study that we describe here. Any deviations from these standard parameters and reasons for those deviations are noted in the respective methods sections.

Energy minimization was performed using steepest descent to a force tolerance of *<* 50 kJ/mol/nm and a step size of 2 pm. Equilibration and production simulations were done with Langevin dynamics using the `sd` integrator with a friction coefficient of 0.2 ps^-1^. The center of mass translational velocity was removed every 10 steps. Lennard-Jones potentials were shifted and cut-off at 1.1 nm, consistent with our previous work (28). We chose a starting value of 1 × 10^−7^ for the `verlet-buffer-tolerance` (units of kJ/mol/ps), a GROMACS-specific setting for Verlet list updates. Setting this parameter correctly is important for reproducibility because using default larger numbers for `verlet-buffer-tolerance` results in simulation instability, due apparently to density inhomogeneity in phase-separated simulation boxes. We found, however, that the `verlet-buffer-tolerance` needed to be altered for proteins with long linkers and stiff linkers. The values of `verlet-buffer-tolerance` that are not the starting value chosen above are documented in Tables S4 and S5. Random seeds for each simulation were generated pseudo-randomly by GROMACS, and were unique for each simulation replicate. Periodic boundary conditions were applied in all three dimensions.

### Simulating LLPS using the slab simulation protocol

We used the slab simulation method from our previous work (28), inspired by other studies (48, 49, 59), to simulate LLPS of CC protein models. We first generate suitable configurations for a protein of interest and pack copies of those configurations into a simulation box to prepare for slab simulation. The slab simulation protocol then consists of the following steps: equilibrating the packed box, compressing the box in one dimension in the NPT ensemble to generate the slab, expanding the box in the same compressed dimension in the NVT ensemble to equilibrate to the desired temperature, and finally a production simulation in the NVT ensemble. For all slab simulations we fix the total number of coil segments in a box to 450 coils, instead of fixing the total number of protein copies, consistent with our previous work (28). This allows us to make comparisons of LLPS propensity between simulations of different proteins.

#### Generating protein configurations by performing single molecule annealing simulations

We start by calculating the number of protein copies necessary to reach our desired total-coil-per-box coil density of 450 coils (Table S2), which is the same density used in our previous study (28). We then prepared that number of unique single molecule simulations to generate the starting configurations that will be packed into a slab simulation box, e.g. preparing 75 single molecule simulations of a 6-coil protein with trimer-forming coils. The starting 3-dimensional CG protein model is generated following the `PeptideBuilder` strategy, discussed in Supporting Material section S.III. The initial CG protein model is then subjected to an annealing single-molecule simulation to generate a configuration that will be packed for the slab protocol. The annealing procedure (described in Supporting Material sections S.IV and S.IV) simulates a protein copy at high temperature (400 K) to disfavor intra-chain coil contacts but allows each segment to relax according to its backbone equilibrium values, and then anneals the protein copy at the desired temperature (e.g. 298 K). Preparing single molecules by annealing allows the protein to reach a low (but not necessarily lowest) energy configuration and allows for the random formation of intra-chain coil contacts. We prepare *N* number of single molecule simulations (Table S2) for each slab replicate to diversify the configurations used for the slab simulation and to mitigate bias that might arise if we picked multiple configurations from one single molecule simulation trajectory. Each slab replicate has its own unique single molecule annealing simulations, e.g. a 6-coil trimer-forming protein will have 225 total individual single molecule simulations (75 configurations per slab × 3 slab replicates).

#### Packing boxes for the slab protocol

Protein configurations, taken from the final frame of single molecule simulations, were used to pack simulation boxes for the slab protocol to ensure that the desired 450 coil segments per box was achieved (Table S2). Slab simulation boxes were made at each desired temperature using single molecule configurations only from the same desired temperature. `packmol` v20.11.1 (60) was used to pack all the starting slab boxes, with a tolerance of 1.0 nm between every protein copy. The box sizes used with `packmol` varied based on the type of protein being packed, and did not represent the final box size used during slab simulation. Proteins with short and 2x-long disordered linkers were packed into 40× 40 ×200 nm boxes, proteins with long disordered linkers were placed into 80 × 80 × 500 nm boxes, and proteins with short stiff linkers were placed into 40 × 40 × 400 nm boxes. We used different starting box sizes to accommodate all the configurations at the specified distance between each copy during packing. Individual protein copies are placed randomly throughout the entire space of the box. Slab simulation boxes were made in triplicate, using only unique single molecule simulation configurations to populate the box.

#### Slab simulation protocol

Packed boxes were energy minimized and then equilibrated in the NPT ensemble for 200 ns with a time step of 20 fs at 150 K, which results in the protein slab used to assess phase coexistence. Table S5 lists proteins that use different slab MD parameters than the default. Semi-isotropic pressure coupling was applied in the *z*-dimension. The Parrinello-Rahman barostat was used with a reference pressure of 1 bar, compressibility of 3 × 10^−4^ bar^−1^, and a time constant of 5 ps for the coupling. We found that using the Parinello-Rahman barostat for 6-coil and 12-coil proteins with 4x-long linkers led to simulation instability. NPT simulations of 6-coil and 12-coil proteins with 4x-linkers used the Berendsen barostat also with reference pressure of 1 bar, compressibility of 3 × 10^−4^ bar^−1^, and a time constant of 5 ps for the coupling. Simulation boxes are compressed to approximately 10-20% of the starting *z*-axis length by the time NPT equilibration finishes.

Proteins across a periodic boundary were then “unwrapped”, and the simulation box was expanded in the *z*-dimension to their final desired length while the slab was kept centered in the box. Proteins with short linkers (disordered, and at various stiffness levels) and proteins with 2x-long linkers had a final *z*-dimension of 200 nm, while proteins with 4x-long linkers had a final *z*-dimension of 300 nm. We used a slightly larger box size for proteins with 4x-long linkers to accommodate their increased size while maintaining a consistent number of total coils in the simulation box as with other proteins. Slabs in extended boxes were then equilibrated at their respective temperature in the NVT ensemble for 200 ns. Production simulations in the NVT ensemble were performed for a total of 20 *μ*s, with the GROMACS `mdrun` parameter `rdd` set to 1.6 nm which is important for domain decomposition. Some protein models used different time steps than the default in the NVT equilibration and production simulation steps of the slab protocol and are detailed in Table S5.

Slab trajectories are modified, prior to any analysis, so that the largest cluster in each frame is centered in the simulation box. Centering the trajectory is important for determining density equilibrium, generating density profiles, and for our custom analyses with python package `mdtraj` (61). We follow a similar trajectory centering algorithm in our previous paper (28). Trajectories are first abbreviated so that only frames in multiples of 10000 ps were used for analysis. Protein molecules were then corrected for movement across periodic boundaries using two sequential calls to `gmx_trjconv`, first using `‘-pbc nojump’` and then using `‘-pbc whole’`. Trajectory centering was then performed in two steps on a frame-by-frame basis. In step 1, coarse density profiles were calculated using `gmx_density` on the periodic boundary-corrected trajectories by dividing the simulation box into twenty equal-sized “slices” along the *z*-dimension. The entire system is then translated so that the slice with the highest density, which should be close to the largest cluster, is approximately centered. In step 2, we then identified the largest cluster in the same frame using `gmx_clustsize` and translated the entire system again such that the center of mass of the cluster is placed in the center of the simulation box. This two step method is necessary to handle the rare instances when in an uncorrected trajectory a slab exists across the periodic boundary in the *z*-dimension, which causes step 2 to fail.

#### Determining equilibrium in slab simulations

We assessed equilibrium by monitoring for number density stability in several locations in the simulation box during the entire course of a slab simulation, similar to previous methods (28). We calculated the average number density in the center of the box with a window of ± 10% of the *z*-axis box length on either side of the center. This region monitors the stability of the putative dense regime. We additionally calculated the average number density across the periodically bound ends of the box, corresponding to the first 12% and last 12% of the *z*-axis box length, to monitor the stability of the dilute region. Number density is calculated per frame using `gmx_density`. The simulation is considered equilibrated when the average density of both the dense and dilute regions is stable. An example density equilibration plots for the 4x-long linker proteins, from one set of slab replicates, are shown in Figure S17. Only equilibrated portions of the trajectories were used for downstream analyses to determine LLPS propensity. All simulations equilibrated within 5–10 *μ*s, consistent with equilibration times seen in our previous study (28).

#### Assessing LLPS propensity in slab simulations

We determine LLPS propensity in slab simulations using both density profile curves and molecular cluster size distributions (Supporting Methods sec. S.VI and S.VII). A phase-separated state must meet the two conditions: (1) an apparent density transition between the dilute and dense regimes of the simulation box, where a transition is defined as ≥ 0.1 number density/nm^3^; and (2) a corresponding molecule cluster that contains ≥ half the number of protein molecules in the simulation.

## RESULTS

### Nonspecifically interacting model coiled-coil proteins can LLPS

We start by establishing the baseline LLPS propensity of CC proteins with nonspecific interactions in order to test hypotheses about how nonspecifically interacting proteins can LLPS. We were motivated by large centrosomal proteins—such as pericentrin, spd-5, and centrosomin, all of which are predicted to contain many individual CC domains—to study model proteins with up to 18 coils per protein.

We observed no phase separation in our initial simulations of nonspecifically-interacting proteins with dimer-forming coils (Fig. 2A); some proteins with trimer-forming coils, however, did phase separate only at the lowest tested temperature (Fig. 2C). Complete density profile curves and molecular cluster sizes analyses are provided in Supporting Figures S9–S12. Simulation snapshots of example dimer-forming (Fig. 2B) and trimer-forming (Fig. 2D) proteins illustrate what LLPS condensates look like for the nonspecifically interacting protein models. The 18-coil trimer-forming proteins show visual evidence of LLPS but suggest that nonspecifically-interacting proteins might be only marginally phase-separating. The overall low LLPS propensity of nonspecifically interacting CC proteins is in contrast to what we found for specific interactions (28), where dimer- and trimer-forming coils readily phase-separated with as few as three coils per protein. We therefore sought to determine what distinguishes the LLPS propensity between trimer- and dimer-forming coil proteins.

**Figure 2.**
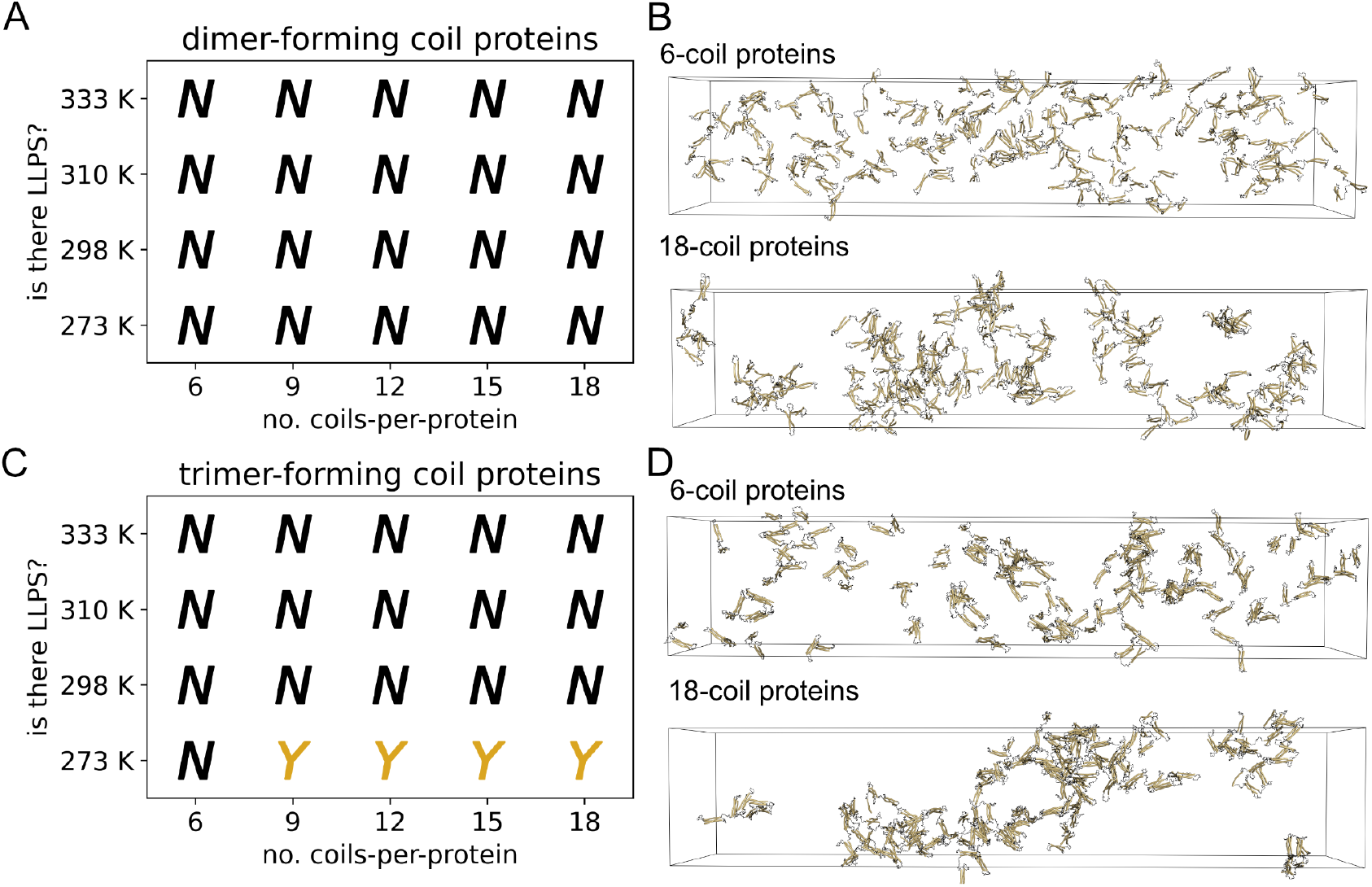
Nonspecifically interacting coil proteins are capable of LLPS. (A) Phase diagram of dimer-forming coil proteins. (B) Example snapshots from slab simulations of 6-coil and 18-coil proteins with dimer-forming coils. (C) Phase diagram of trimer-forming coil proteins. (D) Example snapshots from slab simulations of 6-coil and 18-coil proteins with trimer-forming coils. LLPS assessments made from three replicate slab simulations. Snapshots are from 20 *μ*s at 273 K. Proteins are made whole for visualization, but would wrap through the box boundaries during actual simulation. Visualizations produced using open-source PyMOL v.2.5.0.

It is expected that intra-chain interactions can interfere with LLPS propensity (9) by reducing the number of effective stickers. We hypothesized that we would find a higher propensity for inter-chain contacts in systems that do LLPS compared to systems at similar conditions but which do not LLPS. We found that individual proteins with trimer-forming coils have more available binding sites for inter-chain contacts compared to proteins with dimer-forming coils (Fig. 3). We analyzed individual protein copies prior to starting slab simulations (see Supporting Methods sec. S.V) to determine the capacity for protein copies to form inter-chain coil contacts at the start of the simulation. For each protein, we quantified the number of available binding sites, which is the number of hypothetical inter-chain coils that would be needed to satisfy all unsaturated intra-chain multimers for any given protein configuration. Supporting Figure S2 shows a schematic of example coil multimers and associated available binding sites to illustrate what the metric can physically represent. Increasing polymeric multivalency (i.e. lengthening the protein) does not seem to improve the number of available binding sites for dimer-forming coil proteins, whereas it does seem to increase monotonically with trimer-forming coils. Interestingly, the 9-coil through 18-coil trimer-forming proteins all LLPS with the same relative propensity despite an increasing number of available binding sites.

**Figure 3.**
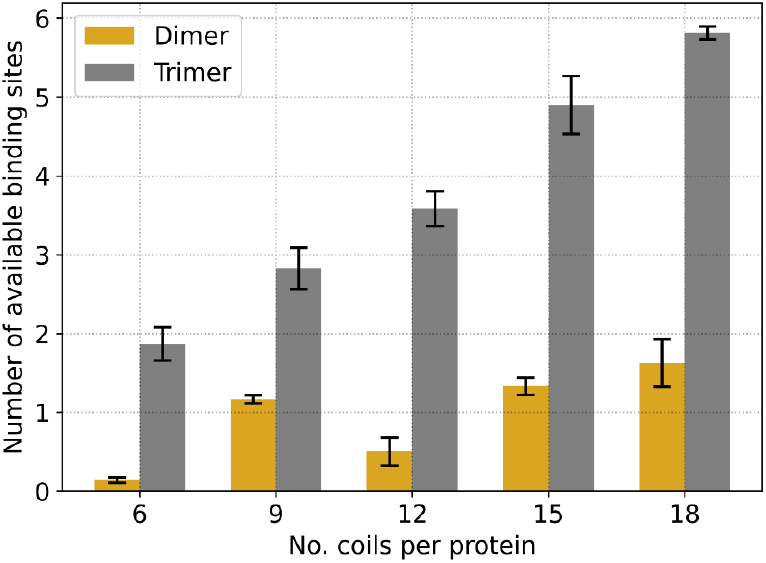
Trimer-forming coil proteins have, on average, more available sites for inter-chain contacts than dimer-forming coil proteins. Plot of number of available binding sites from individual protein configurations at 273 K for each of the types of proteins shown. The number of available binding sites is first calculated and averaged over all the configurations used to pack a single slab replicate simulation (with numbers according to Table S2). The solid bars represent the average of the replicate averaged number of available binding sites, across three slab simulation replicates. Black error bars represent the standard deviation from three slab simulation replicates.

We then examined how intra-chain coil contacts evolve over the course of slab simulations, to explore whether their prevalence might be related to LLPS propensity. We classified the type of contacts that each coil can make—intra-chain, inter-chain, or no contact—and quantified these contact types through the course of each slab simulation (see Supporting Methods sec. S.VIII). Supporting Figures S13 and S14 show that at 273 K for each tested protein showing LLPS, the fraction of intra-chain coil contacts dramatically outnumbers the fraction of inter-chain interactions. Trimer-forming proteins (Fig. S14) demonstrated a fraction of inter-chain contacts that appreciably exceeded the number of unbound coils, which was not true for dimer-forming proteins (Fig. S14). The overall fraction of inter-chain contacts in trimer-forming coils, however, does not completely explain LLPS propensity, as the 6-coil protein does not LLPS yet has approximately the same fraction of inter-chain contacts over time as the other, phase-separating proteins (Fig. S14).

The observed LLPS propensity (Fig. 2) and the predominance of intra-chain coil contacts of tested proteins (Fig. S13 and S14) raised the question: is low LLPS propensity a generic feature of nonspecifically interacting CC proteins? We examined this question by exploring ways in which we might be able to change the ratio of intra-to inter-chain coil contacts, and also LLPS propensity, by changing structural features of the proteins but without changing the intrinsic nonspecific interaction nature of the coil segments.

### Increasing the distance between coil segments has a complex effect on LLPS propensity

We tested the effect of two different linker lengths on the LLPS propensity of 6-coil and 12-coil trimer-forming coil proteins. We used linker lengths 2-times (2x) and 4-times (4x) the length of linkers of proteins in Figure 2 (Supporting Methods sec. S.III). Our rationale for testing longer linkers is that linker size could disfavor intra-chain contacts by spatially sequestering coil segments from each other. Theoretical analyses of associative polymers, in contrast, argues that the excluded volume of the longer chains will actually disfavor LLPS (10). Similarly, recent simulation work from Harmon et al. (17) also suggests that long linkers, evaluated in terms of effective solvation, can inhibit LLPS. We wanted to explore the effect of long linkers on the LLPS of our nonspecifically interacting CC proteins and test if long linkers negatively impact LLPS as others have demonstrated (10, 17).

We found that proteins with 2x linkers have a higher LLPS propensity compared to the 1x-linker proteins (Fig. 4). The 2x-linker proteins form droplets at 273 K that are similar in density to the 1x-linker proteins for both 6-coil (Fig. S15A-B) and 12-coil (Fig. S16C-D) proteins. 4x-linkers show a differential LLPS propensity, where 6-coil proteins do not LLPS but 12-coil proteins can at the same temperatures as the 2x-linker proteins.(Fig. 4A, C). Equilibration analysis for the 4x-linker proteins shows that the 6- and 12-coil proteins are apparently equilibrated (Fig. S17), suggesting that the difference in LLPS propensity is not due to an insufficient simulation time but instead due to effects that the linkers have on the system.

**Figure 4.**
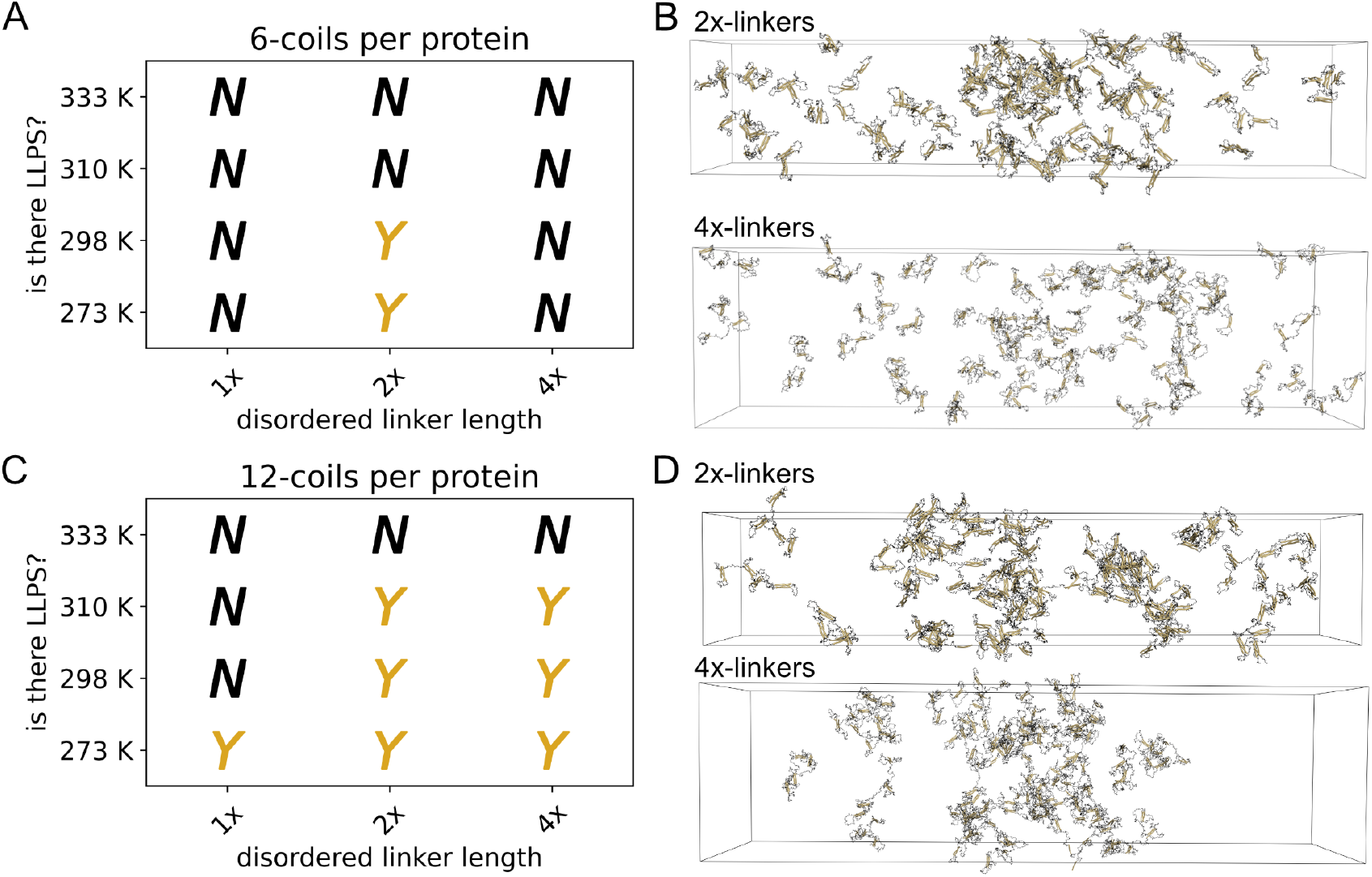
Increasing the linker length has a nontrivial effect on LLPS propensity for proteins with trimer-forming coils. (A) Phase diagram of 6-coil proteins, with trimer-forming coils, at various linker lengths. (B) Example snapshots from slab simulations of 6-coil proteins with 2x- and 4x-long linkers. (C) Phase diagram of 12-coil proteins, with trimer-forming coils, and various linker lengths. (D) Example snapshots from slab simulations of 12-coil proteins with 2x- and 4x-long linkers. LLPS assessments were made from three replicate slab simulations. Snapshots are from 20 *μ*s at 273 K. Proteins are made whole for visualization, but would wrap through the box boundaries during actual simulation. Visualizations produced using open-source PyMOL v.2.5.0.

Increasing linker length seems to increase the fraction of coils that make inter-chain contacts during a slab simulation, though only marginally (Fig. S18). The effect that long-linkers have on increasing inter-chain contacts is most evident for the 6-coil proteins compared to the 12-coil proteins. Interestingly, the 4x-linker length 6-coil proteins show a higher fraction of inter-chain coil contacts than the 1x-linker length analogs (Fig. S18A), but do not have LLPS propensity at any temperature as indicated by profile density and molecular cluster analysis.(Fig. S15).

### Inflexible linker segments strongly enhance LLPS propensity

We hypothesized that stiff linkers separating coil segments will disfavor intra-chain coil contacts and result in LLPS, presumably by reducing nearest neighbor coil contacts along a protein chain because of a high energetic barrier to bending. In previous sections, all the linker segments have been modeled as fully disordered regions (Fig. S19A). Stiff linkers were implemented in our framework by altering the backbone torsional force constant to tune the linker from a disordered into a helical structure. We maintained flexibility at the interface between coil and linker segments by incorporating “hinge” angles/torsions so that different segments could rotate independently relative to each other, and prevent the formation of a long single helix (additional details provided in Supporting Methods sec. S.I and Figure S1). Our experimental collaborators implemented rigid linkers as long single alpha helices in designer CC proteins (26) where they observed improved LLPS propensity with stiff linkers, which motivates our modeling of linkers in a similar way. Importantly, our implementation provides a single tunable parameter to test various degrees of linker stiffness and their effect on LLPS propensity, which would be challenging to explore *in vitro*.

We find that stiff linker segments strongly increase LLPS propensity for both dimer- and trimer-forming coil proteins (Fig. 5A, C). We used three different stiffness levels (presented as 0.*N* relative to coil segment parameters): weak (0.1×, Fig. S19B), moderate (0.5×, Fig. S19C), and strong (0.75×, Fig. S19D). Linkers that are moderately and strongly stiff improve LLPS propensity for both dimer- and trimer-forming coils (Fig. 5A, C). Improved LLPS propensity is also clearly reflected in simulation snapshots for dimer-(Fig. 5B) and trimer-forming (Fig. 5D) proteins. Density profile curves and molecular cluster size distributions for the stiff linker simulations further show a dramatic shift in LLPS propensity as linker stiffness is increased (Figs. S20, S21). The weakly stiff linker proteins did not LLPS at all, for either dimer- or trimer-forming coils.

**Figure 5.**
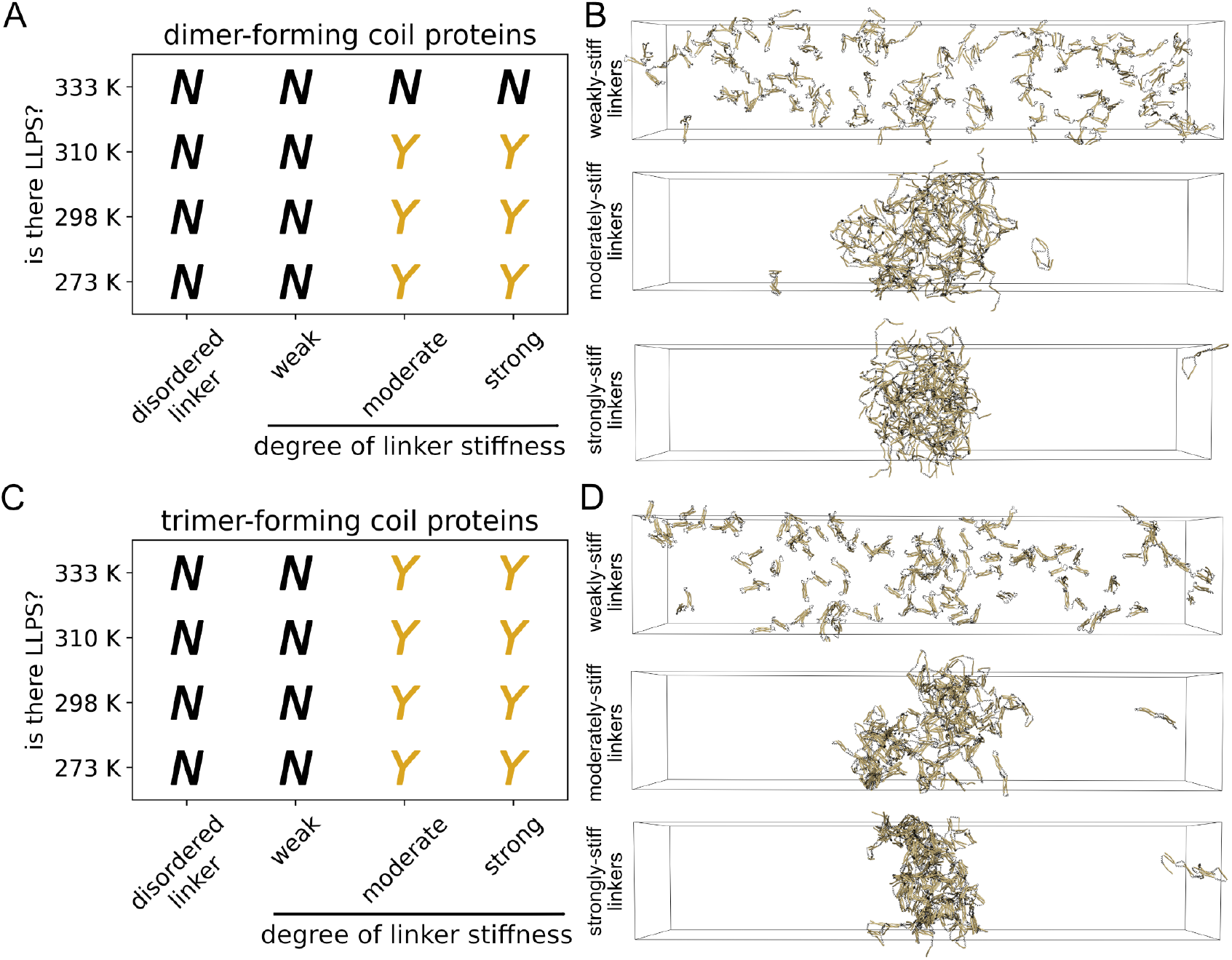
Stiff linkers have a strong effect on promoting LLPS propensity. (A) Phase diagram of 6-coil proteins with dimer-forming coils, with varying degrees of linker stiffness (denoted along *x*-axis). (B) Example snapshots from slab simulations of 6-coil dimer-forming proteins with varying linker stiffness. (C) Phase diagram of 6-coil proteins with trimer-forming coils, with varying degrees of linker stiffness (denoted along *x*-axis). (D) Example snapshots from slab simulations of 6-coil trimer-forming proteins with varying linker stiffness. LLPS assessments were made from three replicate slab simulations. Snapshots are from 20 *μ*s at 273 K. Proteins are made whole for visualization, but would wrap through the box boundaries during actual simulation. Visualizations produced using open-source PyMOL v.2.5.0.

Moderately- and strongly-stiff linkers strongly increase the fraction of inter-chain interactions both for dimer-(Fig. S22A) and trimer-forming coil proteins (Fig. S22B). Proteins with weakly-stiff linkers, in contrast, appeared similar to the disordered linker proteins in terms of coil contact analysis. A striking observation from the coil contact analysis in Figure S22 is the pronounced effect that moderately- and strongly-stiff linkers also have on decreasing the fraction of intra-chain contacts for dimer- and trimer-forming coil proteins. These data largely support our original hypothesis that stiff linkers will disfavor intra-chain interactions, and promote LLPS in nonspecifically interacting coil proteins.

We wanted to understand if the difference in LLPS propensity between weakly- and moderately-stiff linker proteins is a result of the linker structure, or if the weakly-stiff linker simulations are trapped in a metastable state that is affecting our interpretation of results. We performed additional simulations where we swapped the topologies of weakly- and moderately-stiff linker proteins to test if we can take a non-LLPS protein (weakly-stiff) and make it LLPS. The final slab simulation configuration of the weakly-stiff-linker trimer-forming proteins were simulated for an additional 20 *μ*s but with the moderately-stiff linker topologies (i.e. stiffer backbone torsions). Proteins that were originally with weakly-stiff linkers, but swapped to moderately-stiff linkers, were able to form a LLPS-like condensate within the additional simulation time (Fig. S23A-B). Swapping linkers from weakly-to-moderately stiff also decreases intra-chain coil contacts and strongly promotes inter-chain contacts (Fig. S23C) consistent with what is seen for moderately-stiff trimer-forming proteins without the topology-swap (Fig. S22B). The weakly-to-moderately swapped simulations also demonstrate that spontaneous droplet formation can occur within a 20 *μ*s simulation time, since starting with the configuration corresponding to the last frame of the weakly-stiff linker simulations is a dispersed state (Fig. S24) Our results from simulations of proteins with swapped linker stiffness levels demonstrate that the effect of linker stiffness on LLPS propensity is due to the linkers themselves, and not that our simulations are stuck in a metastable state.

### Intra-chain interactions directly affect the LLPS propensity for nonspecifically interacting coiled-coil proteins

We hypothesized that the annealing time for preparing individual protein molecules represents another variable to control the ratio of intra-/inter-chain coil contacts prior to starting slab simulations. By altering the annealing time of our preparative single molecule simulations, we are able to test the sensitivity of our LLPS simulations to initial conditions (i.e. intra-chain coil contacts that exist prior to the start of slab simulations), since we expect the lifetime of individual coil interactions to be relatively long (62). In the previous sets of simulations (Figs. 2–5), we prepared individual protein molecules for slab simulation by allowing each protein copy sufficient simulation time to sample a low energy configurational ensemble through temperature annealing (see Supporting Methods sec. S.IV). We realized that changing the time (shorter than those listed in Table S4) that each protein copy can anneal before the slab simulation starts might change the ratio of preformed intra-chain coil contacts. Examining the radius of gyration plots for individual protein molecules (Figs. S3–S7) shows a short time regime where proteins have not reached a low energy configuration and might correspond to states which leave coils largely unbound and with few intra-chain contacts. Thus, we explored how the annealing time during single protein preparation can affect both intra-chain coil contacts and LLPS propensity of CC proteins without changing the interaction specificity of the coils.

CC proteins prepared with short annealing times readily LLPS (Fig. S25) and do so better than their counterparts from longer annealing times (Fig. 2). Complete density profile curves and molecular cluster size analyses are shown in Figures S26-S27 and S28-S29, respectively. Proteins with trimer-forming coils (Fig. S25B) can LLPS across a much larger range of polymeric multivalencies compared to dimer-forming coils (Fig. S25A), and also compared to their longer-annealing time counterparts counterparts (Fig. 2B).

Trimer-forming coil proteins, generated with short annealing times, also have a high number of available binding sites compared to proteins with dimer-forming coils (Fig. S30) and this corresponds to their increased ability to LLPS. Both proteins with dimer- and trimer-forming coils, from short annealing times, have a greater number of available binding sites prior to slab simulations than those calculated for proteins prepared from long annealing times (Fig. 3). Short-annealed proteins with dimer-forming coils also show a monotonically increasing number of available binding sites as polymeric multivalency is increased (Fig. S30), which was not observed with the long-annealed proteins.

### Mismatched coil lengths do not improve LLPS for nonspecifically interacting proteins

We hypothesized that we could implement preferential coil segment interactions without altering the interaction specificity, and further improve LLPS propensity of proteins prepared with short-annealing times. It is known that the interaction strength between CC domains is dependent on heptad length (63), and that the affinity of CC dimers increases nonlinearly as the number of heptads increase (64, 65). Aronsson et al. showed that *de novo* CC domains of various heptad-lengths could self-sort into heterodimers based on affinities that were seemingly determined by heptad length (66), and we sought to mimic this self-sorting phenomenon. Coil segments that are similar in length could be expected to interact preferentially compared to coils of *mismatched* lengths; this would then reduce the number of possible intra-chain coil contacts without changing the actual specificity of the coil interactions and might result in increased LLPS propensity.

We found, however, that proteins that contain mismatched coil segment lengths (and thus, different number of heptads) do not LLPS better than proteins with consistent coil segment lengths prepared with short-annealing times (Fig. S31, profile density and molecular cluster distribution plots in Figures S32-S36). In each of the plots in Figure S31, the different proteins tested are represented by a string of “C”s and “M”s to highlight where the mismatched coils are, with “C” representing a normal coil with 5-heptads (similar to previous protein models), and “M” representaing a mismatched coil with 3-heptads. We find that some of the mismatched-coil proteins can still LLPS at a similar propensity to the proteins without mismatch, but in most cases introducing the shorter coils results in a reduced LLPS propensity (Fig. S31). We did see that most proteins with mismatched coil segments lengths still had greater LLPS propensity than proteins prepared from long-annealing times (Fig. 2). Even within the short annealing regime, which favors LLPS (Fig. S25), mismatched-coils did not further improve the LLPS propensity of the tested model CC proteins.

## DISCUSSION

We report in this paper: (1) model CC proteins can LLPS with nonspecific interactions between the CC domains (stickers); (2) intra-sticker interactions compete with inter-sticker interactions and can negatively affect LLPS propensity; and (3) linker segments (i.e. spacers) can positively affect LLPS propensity by reducing the formation of intra-coil interactions in model CC proteins. Importantly, our results also demonstrate that LLPS droplets we observe are not just kinetically-arrested metastable states. Figures S23 and S24 show that droplets can form from a fully dispersed set of proteins, indicating that LLPS can be a thermodynamically achievable end-result for our nonspecifically interacting proteins.

### Linkers can affect LLPS propensity by affecting how a protein interacts with itself

Our work and recent studies demonstrate that the effect linkers have on intra-chain interactions seems to be an important determinant for LLPS propensity. Our study shows that long linkers can mostly improve the LLPS propensity of nonspecifically interacting CC proteins (Fig. 4), which is somewhat in contrast with polymer theory that predicts long linkers would disfavor LLPS (10). We additionally found that stiff linkers can greatly enhance LLPS propensity of nonspecific CC proteins (Fig. 5). This result is also in contrast with a previous simulation study that found that linkers which prevent nearest neighbor sticker interactions did not support LLPS (17). We rationalize the difference in our results to previous mean-field studies by focusing less on the exact structure of the linker, and more on the effect that our long and stiff linkers have on changing intra-chain interactions (Fig. S18 and S22) which are necessary for LLPS. Work by Chen et al. (67) and Zhou et al. (68) demonstrate the effect that linkers can have on intra-chain interactions and LLPS propensity by studying the condensation of nucleosome arrays driven by nucleosome particles. They show that individual nucleosome arrays do not LLPS with long DNA linkers, because long DNA linkers promote intra-array interactions between nucleosomes which reduces array multivalency (68). Arrays with short DNA linkers, in contrast, do LLPS because short DNA linkers impose a geometrical torsion restraint on neighboring nucleosomes that prevents intra-array nucleosome contacts (67). These studies (67, 68) are clear examples that correlate how linker size can affect LLPS by promoting or preventing the formation of intra-chain contacts. Our current study, in addition to the work with nucleosome arrays (67, 68), suggests that the effect linkers have on LLPS should be viewed, in part, by how linkers favor inter-versus intra-chain interactions instead of just simply on the length or the structure of the linker.

### LLPS might broadly be a feature of coiled-coil proteins

Our finding that model CC proteins can LLPS with nonspecific interactions has important potential implications for the general LLPS propensity of natural CC proteins. Specific interactions between CC domains are known to be important for the LLPS of some model (28) and designed CC proteins (26, 27). We expect naturally occurring CC domains will interact specifically with their cognate CC partner(s), based on structural specificity between the interacting CC domains (21, 23, 55) conferred by amino acid sequence. It is likely, however, that CC domains and proteins will also form nonspecific interactions with themselves/other biomolecules (40, 69) in cells largely due to the crowded nature (70) of the cytosol and other compartments.

The contributions of nonspecific interactions to LLPS are not well studied, but it is known that nonspecific interactions arising from molecular crowding can seem to promote LLPS (40, 71), and that nonspecific interactions can sometimes help the formation of specific interactions and lead to LLPS in simulated systems (44). Our current work demonstrates that LLPS can still arise from purely nonspecific interactions, and adds to the field’s understanding about these types of interactions and their role in LLPS.

The ability of model CC proteins to LLPS with both specific and nonspecific interactions suggests that CC-driven LLPS might be a generally accessible phenomenon for CC proteins, especially if we consider that natural CC proteins are likely to form both specific and nonspecific interactions simultaneously in a crowded cellular environment. This might be particularly true for CC proteins with multiple CC domains, and/or for proteins whose CC domains can form trimers or higher order multimers, since we show in this and previous work (28) that multivalency supports CC-driven LLPS. It would not be surprising for CC-driven LLPS to be a widespread phenomenon for CC proteins, since it is now expected that *most* proteins are capable of forming LLPS droplets (72, 73), though typical cellular conditions might just result in transient LLPS states (73). It will become more important to focus on the cellular conditions that allow for LLPS of CC proteins—instead of simply asking if LLPS is possible, considering that most proteins probably are capable—so that we can identify biologically relevant LLPS (74).

### Intra-chain interactions might play important regulatory roles in controlling LLPS within cells

Our findings demonstrate that intra-chain interactions can interfere with a protein’s ability to LLPS, e.g. for CC proteins with disordered linkers (Figs. 2, 3, S13, and S14). We further show that if we take the same type of proteins but generate configurations intentionally to have fewer intra-chain interactions, then LLPS propensity broadly increases (Fig. S25), and this occurs without changing the structure or dynamics of linker segments. Our results suggest that changes to a protein’s configurational ensemble is another potential method to modify a protein’s LLPS propensity that doesn’t require changes to the structure or sequence of the protein. Cells might be able to regulate protein LLPS by changing a protein’s configurational ensemble, such as for the *C. elegans* centrosomal protein spd-5. The current model for spd-5 phase separation starts with the protein in an autoinhibited state where spd-5 forms multiple intra-chain interactions, but upon activation and phosphorylation by PLK-1 some intra-chain interactions are disrupted which allows inter-CC interactions to form (34). Environmental conditions, such as salt concentration, pH, crowding, and temperature, will also change the folding equilibrium of proteins and alter their configurational ensemble, which can affect how readily accessible stickers are to other proteins. A review by Scholl and Deniz (75) highlights four key examples of proteins—FUS, NPM1, K18, and ELP—that experience changes in their configurational ensemble that (largely positively) affect their LLPS propensity.

Cells can also use molecular chaperones to affect the LLPS propensity of proteins, such as for the disordered protein FUS. Boczek et al. (76) showed that the chaperone HspB8 can be recruited into FUS droplets and prevent misfolding of the FUS RNA-binding domain, which ultimately keeps FUS from forming amyloid-like structures and maintaining a liquid-phase of the droplet. Molecular chaperones are also thought to affect pathological condensate formation of other ALS-related proteins similar to FUS by helping them avoid amyloid-like structures (77–79). The impact of chaperones on the phase separation of other proteins and how they can specifically change intra-chain interactions to affect LLPS of substrate proteins remains understudied, however. The ability for cells to control protein configurations, and therefore LLPS propensity, in response to changing cellular conditions or through molecular chaperones can be viewed as a consequence of cells existing at nonequilibrium conditions (80).

## SUPPORTING MATERIAL DESCRIPTION

Online Supporting Material for this document can be found on bioRxiv https://www.biorxiv.org/. Document S1 (PDF) contains supporting methods, figures S1–S36, and tables S1–S5.

## ACKNOWLEDGMENTS

This work utilized the Alpine high performance computing resource at the University of Colorado Boulder. Alpine is jointly funded by the University of Colorado Boulder, the University of Colorado Anschutz, Colorado State University, and the National Science Foundation (award ACI-2201538). This work was supported by the NIH Molecular Biophysics Training Program (T32GM065103, D.A.R.), NIH fellowship F31GM151838 (D.A.R.), NSF MCB-1943488 (L.E.H.), and a University of Colorado Research and Innovation Seed Grant (M.R.S., L.E.H.).

## AUTHOR CONTRIBUTIONS

Contributions according to CRediT Taxonomy: **D. A. R**.: Methodology, software, investigation, visualization, writing. **A.S**.: Investigation, software, visualization. **L. E. H**.: Conceptualization, writing, supervision, project administration. **M. R. S**.: Conceptualization, writing, supervision, project administration.

## SUPPORTING CITATIONS

References (81–84) appear in the Supporting Material.

