## Supporting Material for "Nonspecific interactions can lead to liquid-liquid phase separation in coiled-coil proteins models"

**Supporting Material – Document 1**

### SUPPLEMENTAL METHODS

#### S.I Structural parameters of CC LLPS framework

We use the same structural parameters from our previous publication (1). All CG beads have a mass of 109.0 amu, which is approximately the average mass of the 20 proteinaceous amino acids. Bond stretching between each CG bead is modeled with a harmonic potential (equation 1) with an equilibrium distance of 0.385 nm and a force constant of 50000 kJ/mol/nm$^2$.

Pseudo-bond angles (defined between three CG beads) are treated with a harmonic potential (equation 2) with an equilibrium angle of 93 degrees and a force constant of 50.0 kJ/mol/rad$^2$. Pseudo-torsions (defined between four CG beads) are handled by a linear combination of two periodic cosine potentials (equation 3). The first potential uses an equilibrium potential of -130.0 degrees, force constant of 20 kJ/mol, and multiplicity $n = 1$. The second potential uses an equilibrium potential of -30.0 degrees, force constant of 20 kJ/mol, and multiplicity $n = 3$.

The equilibrium values for the pseudo-bond angles and -torsions correspond to a helical structure in the reduced CG space, mapped from the all-atom space (2) and based on the average $\phi$,$\psi$ torsions from crystal structures of alpha helices (1). The above parameters are used exactly for the coil segments and result in a structure that has some bending flexibility in the long axis of the helix but the overall helical shape is preserved. Disordered linker segments use the same equilibrium values angles and torsions as in the coil segments. Linker segments, however, use angle and torsion force constants that are 0.01× that of the coil segments.

$$V_b(r_{ij}) = \frac{1}{2}k_b(r_{ij} - b_{ij})^2 \tag{1}$$

$$V_a(\theta_{ijk}) = \frac{1}{2}k_\theta(\theta_{ijk} - \theta^0_{ijk})^2 \tag{2}$$

$$V_d(\phi_{ijkl}) = k_\phi(1 + \cos(n\phi - \phi_s)) \tag{3}$$

The strength of the pseudo-angles and -torsions in the linker segments represents a variable that can be modified to change the behavior of the linker structure. We generated linker segments of varying stiffness by changing the multiplicative factor that scales the force constants relative to the default coil segment values. We used the following scale factors to design differentially stiff linker segments (which are in addition to the disordered linker segments): 0.1× weaker for “weakly stiff" linkers; 0.5× weaker for “moderately stiff" linkers, and 0.75× weaker for “strongly stiff" linkers.

It was necessary to insert “hinge" pseudo-angles/-torsions between the coil segments and stiff linker segments to allow the segments to move freely relative to each other. Without the hinge pseudo-angles/-torsions, a protein with stiff linkers would end up as a semi-stiff rod along the entire length of the sequence, which is unphysical. We designed two hinge pseudo-angles, shown schematically in Figure S1A, with angle 1 being the last two beads of a segment plus the first bead of the next segment, and angle 2 being the last bead of a segment plus the first two beads of the next segment. Three hinge pseudo-torsions were used, shown in Figure S1B, and are defined such that torsion 1 is the last three beads of a segment plus the first bead of the next segment, torsion 2 is the last two beads of a segment plus the first two beads of the next segment, and torsion 3 is the last bead of a segment plus the first three beads of the next segment. The equilibrium values for these hinge regions are the same as for all other segments, but the force constants are 0.01× that of the coil segment constants.

Each bead has four neighbors excluded from nonbonded interactions. Explicit 1-4 and 1-5 nonbonded pair interactions are used in the both the coil and linker segments and are described with a 12-6 Lenanrd-Jones (LJ) potential (e.g. equation 4). 1-4 pairs interact with $\epsilon = 2$ kJ/mol and $\sigma = 0.453$ nm. 1-5 pairs interact with $\epsilon = 2$ kJ/mol and $\sigma = 0.561$ nm. Linker segment beads use the same sigmas, but with scaled epsilon values that match the same level of scaling used for the pseudo-angles/-torsions (i.e. weakly stiff linkers have 0.1× weaker 1-4/1-5 epsilons). 1-4 and 1-5 pair interactions do not cross between coil and linker segments.

#### S.II Nonspecific interaction parameters of CC LLPS framework

We use the same classes of bead types in the present study as we used in our previous work (1). The three bead classes in our framework include: *coil-coil sticky beads*, *multimer-driving beads*, and *repulsive-only beads*. Coil-coil sticky beads exist only in coil segments, drive the association between coil segments, and are placed in positions *‘a’* and *‘d’* of the heptad repeat. Multimer-driving beads help control the type of multimer the coil segments can form and are in positions *‘e’* and *‘g’* of the heptad repeat. Repulsive-only beads correspond to beads that provide no net-attractive protein–protein interactions and are used in positions *‘b’*, *‘c’*, and *‘e’* of the heptad repeat. Repulsive-only beads also make up the entirety of linker segments.

GROMACS allows the use of a simplified 12-6 Lennard Jones potential as follows:

$$V_{LJ}(r_{rj}) = \frac{C_{12,ij}}{r_{ij}^{12}} - \frac{C_{6,ij}}{r_{ij}^{6}} \tag{4}$$

where,

$$C_{12,ij} = 4\epsilon_{ij}\sigma_{ij}^{12} \tag{5}$$

and,

$$C_{6,ij} = 4\epsilon_{ij}\sigma_{ij}^{6} \tag{6}$$

The coil-coil sticky beads and the multimer-driving beads utilize both the $C_{12,ij}$ and $C_{6,ij}$ terms. The repulsive-only beads, in contrast, use only the $C_{12,ij}$ term (therefore setting the attractive component equal to zero) to model only repulsive interactions. Table S1 lists all the interaction terms used in the framework and is modified from the table published in Ramirez et al. 2024 (1) since the interaction values remain the same between this study and the previous one. All of the bead interaction terms were parameterized to reflect the behavior of CC domains and disordered domains in solution, which allows us to use this framework with the solvent treated implicitly. No explicit Coulombic interaction terms were used for this study.

#### S.III Generating coarse-grained coiled-coil single proteins via the `PeptideBuilder` strategy

CG CC proteins for use in our framework are constructed with the python package `PeptideBuilder` v1.1.0 (3). Initial protein models constructed with `PeptideBuilder` are represented atomistically, and are C-$\alpha$ coarse-grained using the python package `ProDy` v2.3.1 (4, 5). This approach is identical to the method we used previously for generating coarse-grained proteins (1). The default length of coil and linker segments is 35 (i.e. 5 heptads) and 25 CG beads, respectively, similar to our previous work. The segment lengths can be changed at will, however, such as to generate 2x-long (50 CG beads) and 4x-long linkers (100 CG beads), or to change the coil segments to three heptads (21 CG beads). Protein configurations generated via PeptideBuilder put both the coil and linker segments in helical geometry regardless of the desired geometry specified in the GROMACS toplogy file. Single molecule MD simulations are thus necessary to relax the protein segments into their equilibrium values.

#### S.IV Annealing simulation procedure for preparing single protein molecules

Proteins generated using the `PeptideBuilder` strategy (Supporting Material sec. S.III) were subjected to a single molecule annealing simulation to equilibrate the structure and prepare it for slab simulation.

The generated single protein copy was placed into a cubic simulation box large enough to accommodate the initial structure plus ~10 nm on all box vectors. The protein was energy minimized, and then simulated in the NVT ensemble at 400 K (the “hot step") for the corresponding time listed in Table S3, using either the default 25 fs or a time step indicated in Table S4. Proteins were then annealed in the NVT ensemble at the desired slab simulation temperature (the “annealing step") according to the times listed in Table S4, again using either the default 25 fs time step or a time step indicated in Table S4.

Immediately following the end of the annealing simulation step, the trajectories are converted using `gmx_trjconv` so that the protein copy is made whole (and is not broken across a periodic box boundary) and centered in the box for every frame. The configuration from the final frame is saved and used as input for packing boxes for slab simulations. The annealing step trajectories are analyzed for radius of gyration using the GROMACS utility `gmx_gyrate`, as well as the number of available binding sites.

##### Confirming the appropriate annealing time for single molecule simulations

Our goals for the single molecule annealing simulations were to have a protein copy (1) relax into its equilibrium backbone angles, and (2) reach a relatively low energy configuration, and so we selected single molecule annealing times that would allow us to achieve these goals. We did not expect that one single-molecule annealing simulation time would be appropriate for all the types of proteins studied—disordered linkers, long linkers, and stiff linkers—due to differences in total protein length/flexibility affecting protein dynamics and sampling, and so we varied the annealing time for the different proteins studied. We used the radius of gyration ($R_g$) and the number of available binding sites (next section) as the metrics to assess that our protein configurations have reached a low energy configuration. Both the $R_g$ and number of available binding sites reaching a stable average over time would suggest the protein has reached a low energy configuration and indicate that the single molecule annealing simulation time was appropriate.

We analyzed and averaged the $R_g$ timeseries of the entire annealing step across all the single molecule simulations, at 273 K, belonging to one slab replicate. Three averaged $R_g$ timeseries, corresponding to three slab replicates, were then averaged again to produce a single $R_g$ timeseries for a given protein at 273 K. The $R_g$ from the single molecule simulations shows that the metric stabilized within the annealing simulation time for disordered linkers (Fig. S3), long linkers (Fig. S5), and stiff linkers (Fig. S7), indicating that each type of protein can sample from a low energy ensemble within the allotted annealing step. The number of available binding sites throughout the annealing simulation was also calculated and averaged across each single molecule simulation at 273 K (description in Supporting Methods S.V). The averaged available binding sites timeseries was then averaged again across three slab replicates' worth of single molecule simulations for a given protein. Analysis of number of available binding sites over time also stabilize for most of the the different types of proteins (disordered linkers, Fig. S4; long linkers, Fig. S6; stiff linkers, Fig. S8). Available binding site analysis for proteins with disordered linkers and 18-coils do not appear to be entirely converged, but are clearly close to such a state (Fig. S4). Only the 6-coil dimer-forming disordered linker proteins (Fig. S4) and the 6-coil dimer-forming weakly stiff-linker proteins (Fig. S8) result in a number of available binding sites close to 0 at the end of the annealing simulation step. The other proteins sample from low energy configurations that result in a number of available binding sites > 1. These two analyses suggest that the annealing times for each type of protein (disordered linkers, long linkers, and stiff linkers) is sufficient to generate relatively low energy configurations for slab simulation.

Importantly, the annealing time can be changed at-will to generate protein configurations that sample from higher energy configurational ensembles to intentionally change the amount of intra-chain contacts present, if desired.

#### S.V Calculating the number of available binding sites in protein configurations

We assess the capacity of individual protein models to form inter-chain contacts through quantifying the *number of available binding sites* in a given configuration. This metric identifies unsaturated coil multimers and determines the number of hypothetical inter-chain coils that would be necessary to make the unsaturated multimers become saturated. Supporting Figure S2 demonstrates a physical interpretation of available binding sites in our model CC systems.

The number of available binding sites are calculated using a custom python script that finds both coil multimers and unbound coils in a simulation or a single configuration. The distance between all coils' centers of mass are calculated, following a procedure similar to our previous work (1) and Supporting Method S.VIII, and multimers are identified if coils are within a 1.3 nm cut-off distance of each other. Completely unbound coils and unsaturated multimers are then selected, and the total binding sites for each type identified is calculated based on the type of multimer that the coils can form (Fig. S2).

Importantly, the number of available binding sites analysis can be performed as a timeseries throughout a simulation, or can be calculated on a configuration from a single frame of a simulation. We utilized both ways to help determine if the single molecule annealing simulation times could generate low energy protein configurations (via timeseries, Supporting Methods sec. S.IV), and to assess the ability for protein configurations to form inter-chain interactions prior to starting a slab simulation (via single frame analysis). The code that handles the calculation of available binding sites (see Methods, "Code Availability") is capable of analyzing multiple proteins over time, i.e. a slab simulation, but the output is configured to be the *total* available binding sites for the entire system; we caution against using this analysis method on slab simulation trajectories.

#### S.VI Density profile analysis

We performed density analysis of slab-centered trajectories, only on portions that are at density equilibrium, using the `gmx_density` module in GROMACS. Density (reported as number density / $nm^3$) was calculated in 2 nm slices along the $z$-axis of the simulation box, and the average density in each 2 nm slice is recorded. Density profiles along the $z$-axis were averaged across three slab replicates for each temperature and are plotted as mean (solid lines) ± standard deviation (shaded regions) in all density profile plots. Density profiles that do not correspond to a phase-separated state will still show a tiny bump at the center of the profile due to centering the trajectory of the largest cluster in each frame.

#### S.VII Molecular cluster analysis

Molecule cluster size distributions were generated from the equilibrated and slab-centered trajectories using the `gmx_clustsize` analysis tool in GROMACS. Cluster sizes for whole molecules were calculated using a distance cut-off of 0.9 nm between interaction pairs. The output cluster size populations were normalized to probabilities and averaged across three slab replicates for each of the temperatures. Data are reported as mean ± standard deviation over three replicate simulations

#### S.VIII Identifying types of coil–coil contacts in slab simulations

We identified the types of coil–coil contacts that are made in slab simulations—intra-chain, inter-chain, or unbound—and quantified the fraction of each in a two-step process using custom python scripts. The first step determines coil multimers in desired frames of a simulation by calculating the distance between the center of mass of all coil segments. Coil segments within a 1.3 nm cut-off of other coils are said to be in a multimer, and the type of multimer (dimer, trimer, etc.) is determined.

The second step uses the data from the first step to calculate the types of coil–coil contacts. For each analyzed frame, the script loops through every identified multimer and determines if a coil within the multimer is making an intra-chain interaction, an inter-chain interaction, *or both*. For multimers with three or more coils, there is a possibility that a given coil can make both intra-chain and inter-chain interactions, but our custom script tracks only that a certain type of interaction is formed for a given coil. Thus, each type of coil–coil contact, summed for all coil segments, can be represented as a fraction of the total number of coils. For this reason, however, it is not true that the sum of the fraction of inter-chain, intra-chain, and unbound contacts in a single frame is guaranteed to be $\leq 1$.

#### SUPPLEMENTAL FIGURES

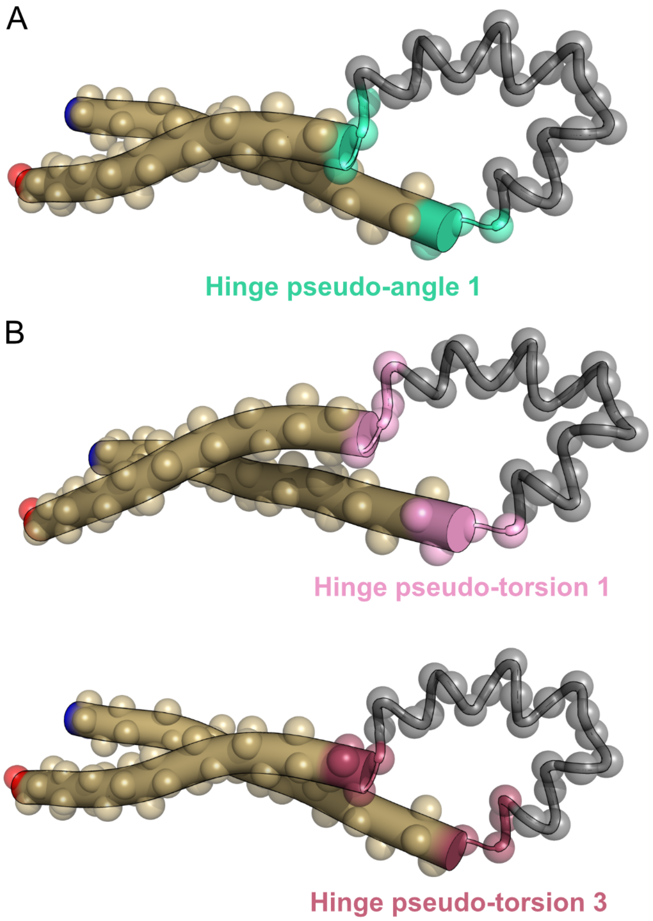

Figure S1: Schematic of the hinge regions that separate coil segments from stiff linker segments. Individual (A) pseudo-angles and (B) pseudo-torsions that define the hinge region for proteins with stiff linkers are shown in colors that match the text and are numbered consistent with the methods text. Solid golden cylinders represent coil segments, and grey tubes represent linker segments. C-$\alpha$ CG beads are also shown as ghosted spheres to illustrate beads that participate in the different hinge angles/torsions. The dark blue and bright red beads in each image highlight the N- and C-terminal beads to help orient the viewer. Images were generated with PyMOL v.2.5.0. on a randomly chosen protein with moderately stiff linkers.

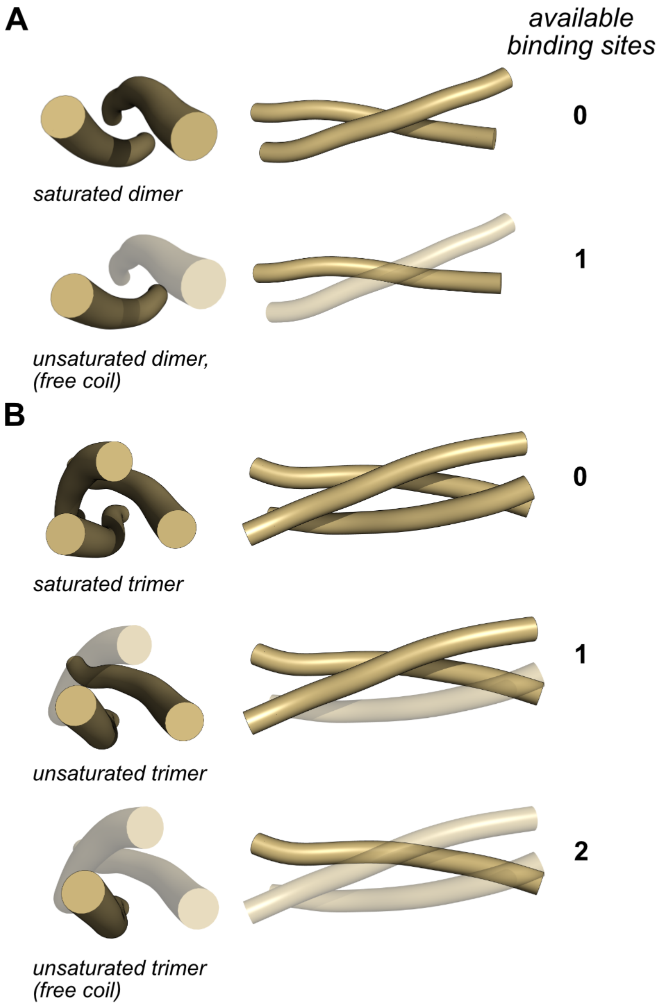

Figure S2: Available binding sites represents a capacity to form inter-chain contacts. Schematic of saturated and unsaturated (A) dimers and (B) trimers to demonstrate the available binding sites metric. Ghosted helices visually show coil vacancies for each type of unsaturated multimer, which physically represents the number of available binding sites i.e. inter-chain coils that would be needed to saturate the multimer.

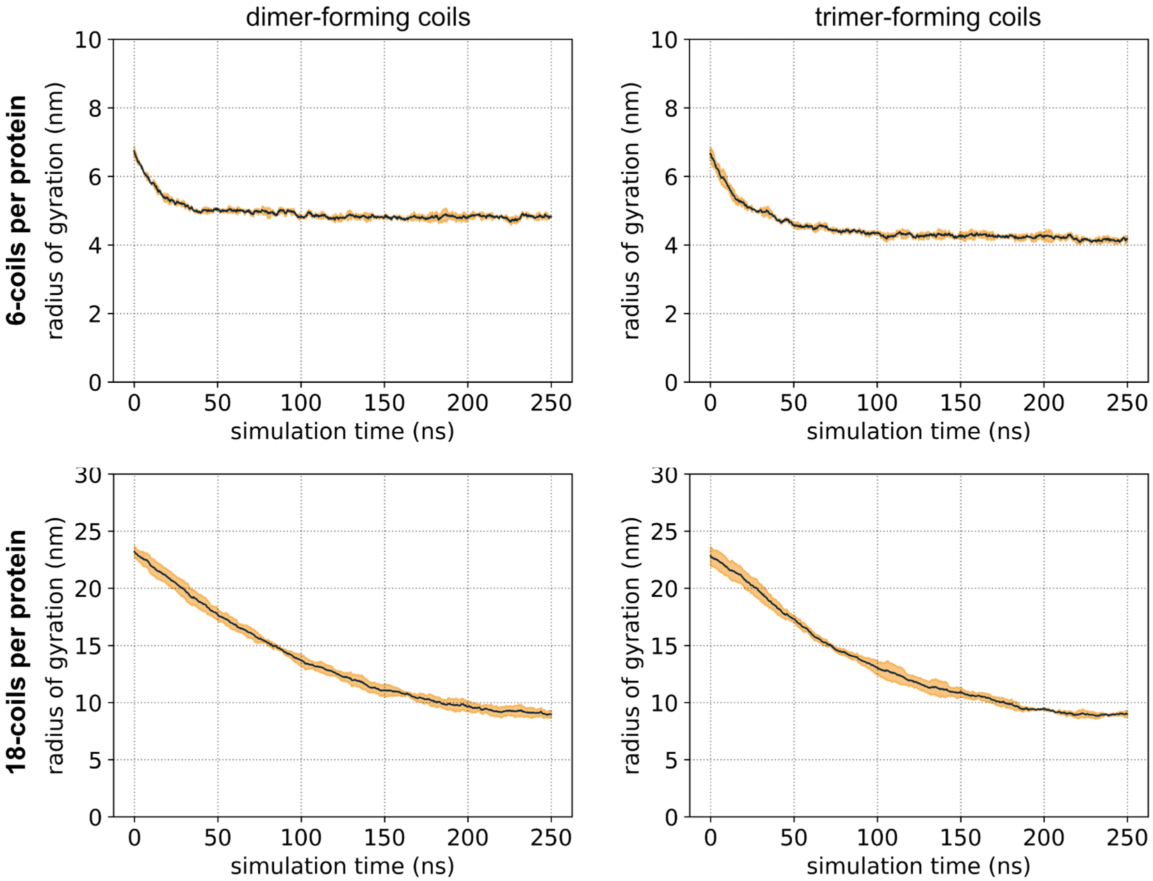

Figure S3: Radius of gyration analysis of single protein molecules during the annealing step of the protein preparation procedure. Data are from single molecule simulations at 273 K only. The radius of gyration of individual prepared protein molecules was averaged over all protein molecules for a slab replicate (*n* according to Table S2), and then the average was averaged across three slab replicates to produce the solid dark blue line. Orange shading around the mean represents the standard deviation of the replicate averaged radius of gyration.

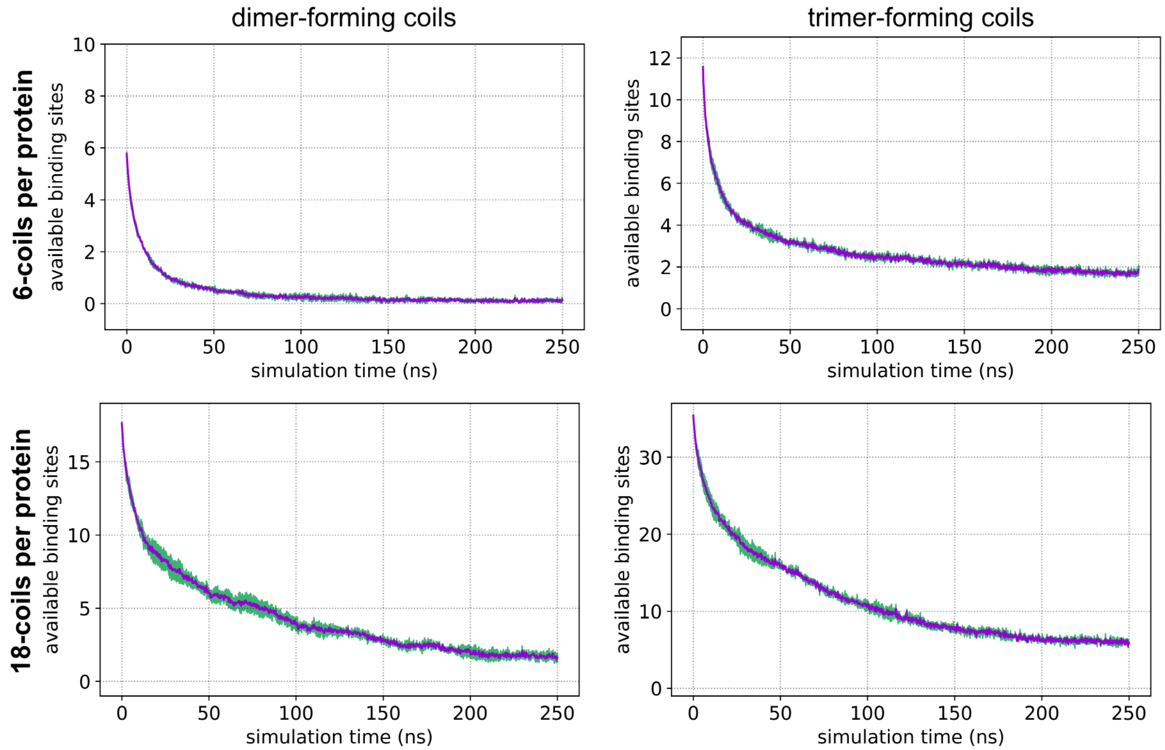

Figure S4: Number of available binding sites of single protein molecules during the annealing step of the protein preparation procedure. Data are from single molecule simulations at 273 K only. The number of available binding sites of individually prepared protein molecules was averaged over all the protein molecules for a slab replicate (*n* according to Table S2), and then the average was averaged again across three slab replicates to produce the solid purple line. Green shading around the mean represents the standard deviation of three slab replicate averaged data.

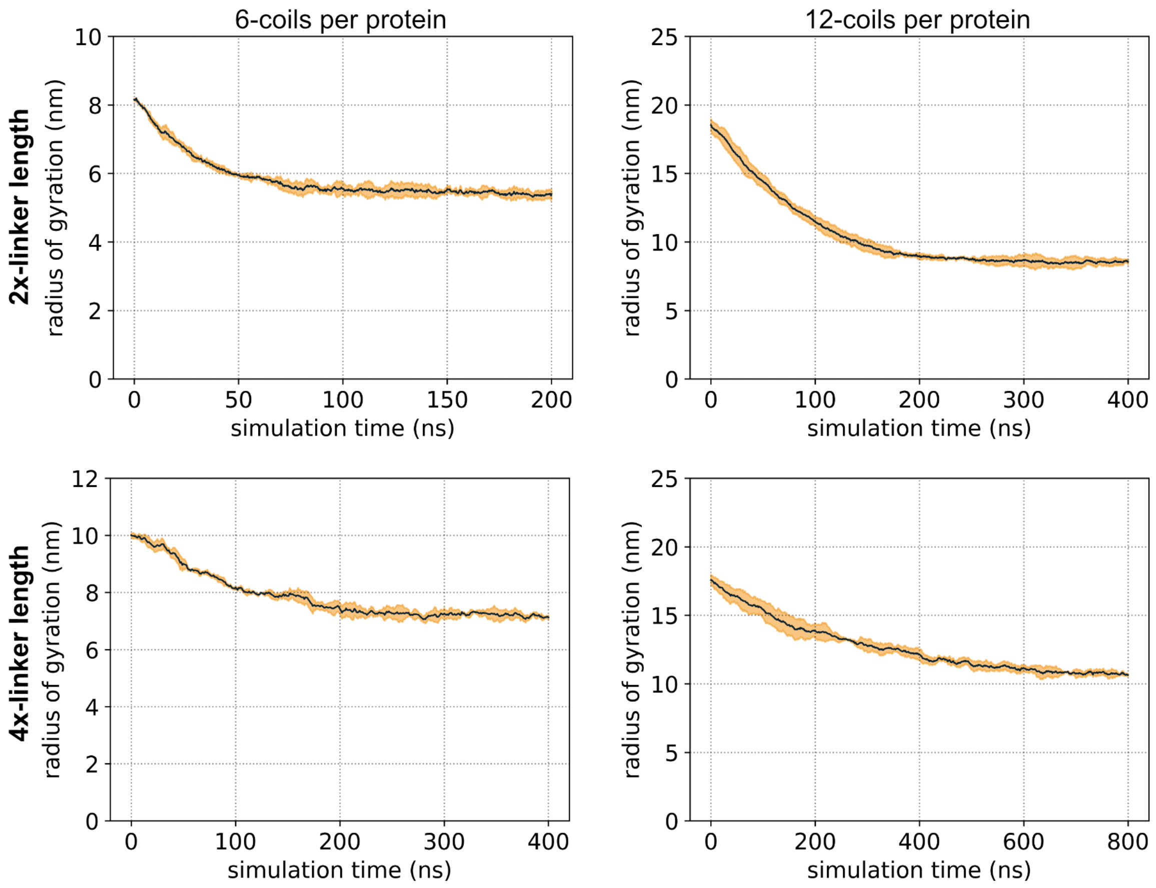

Figure S5: Radius of gyration analysis of single protein molecules with long linkers during the annealing step of the protein preparation procedure. Data are from single molecule simulations at 273 K only. The radius of gyration of individual prepared protein molecules was averaged over all protein molecules for a slab replicate (*n* according to Table S2), and then the average was averaged across three slab replicates to produce the solid dark blue line. Orange shading around the mean represents the standard deviation of the replicate averaged radius of gyration.

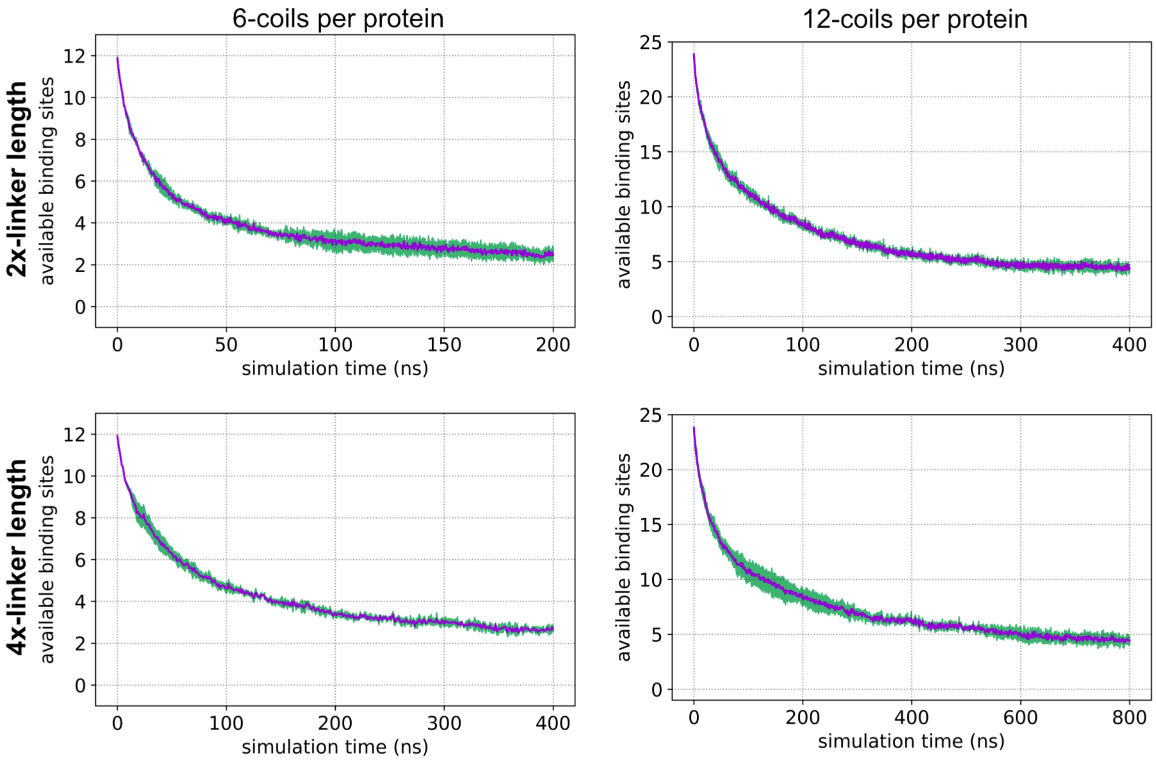

Figure S6: Number of available binding sites of single protein molecules with long linkers during the annealing step of the protein preparation procedure. Data are from single molecule simulations at 273 K only. The number of available binding sites of individually prepared protein molecules was averaged over all the protein molecules for a slab replicate ($n$ according to Table S2), and then the average was averaged again across three slab replicates to produce the solid purple line. Green shading around the mean represents the standard deviation of three slab replicate averaged data.

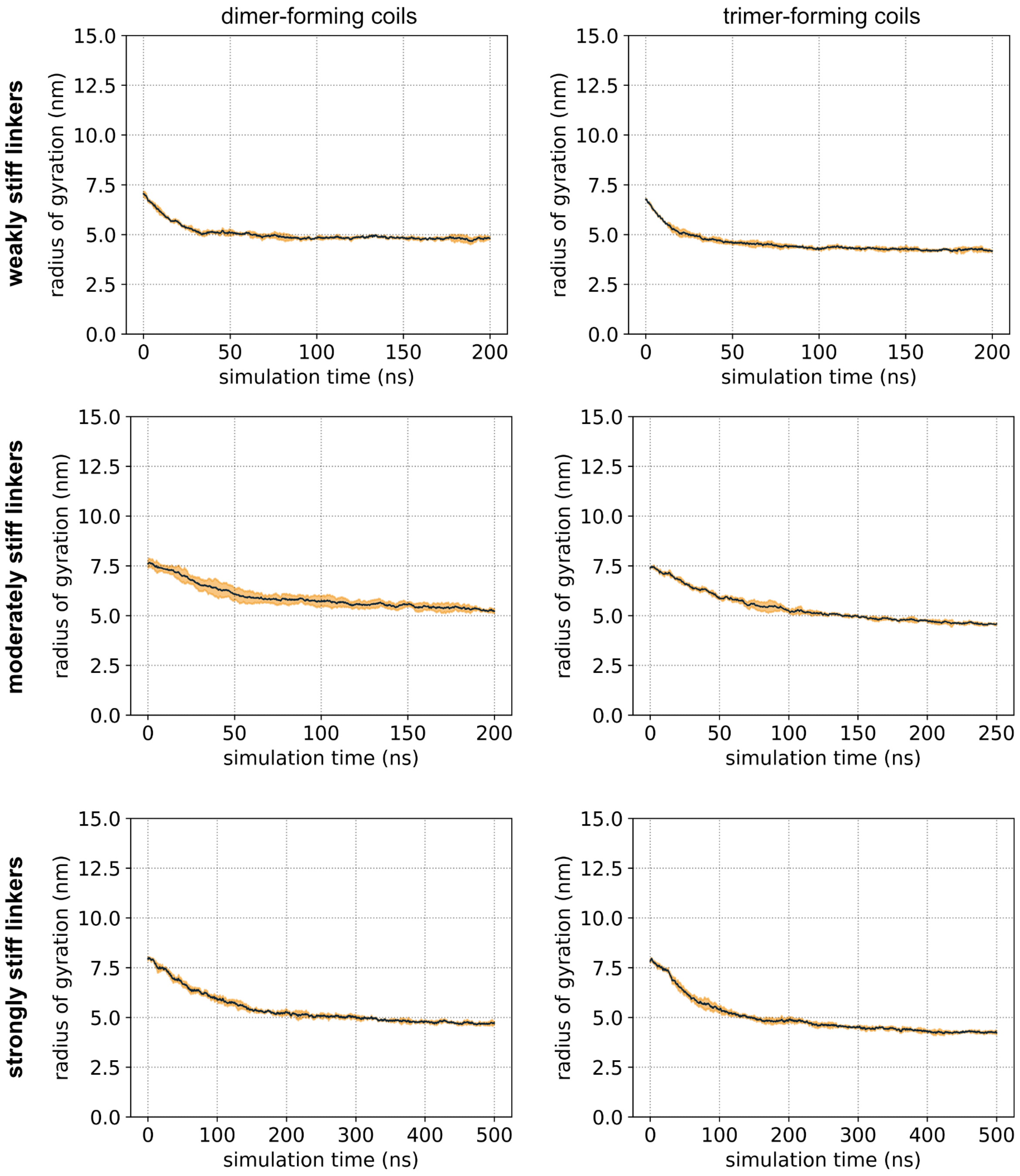

Figure S7: Radius of gyration analysis of single protein molecules with stiff linkers during the annealing step of the protein preparation procedure. Data are from single molecule simulations at 273 K only. The radius of gyration of individual prepared protein molecules was averaged over all protein molecules for a slab replicate (*n* according to Table S2), and then the average was averaged across three slab replicates to produce the solid dark blue line. Orange shading around the mean represents the standard deviation of the replicate averaged radius of gyration.

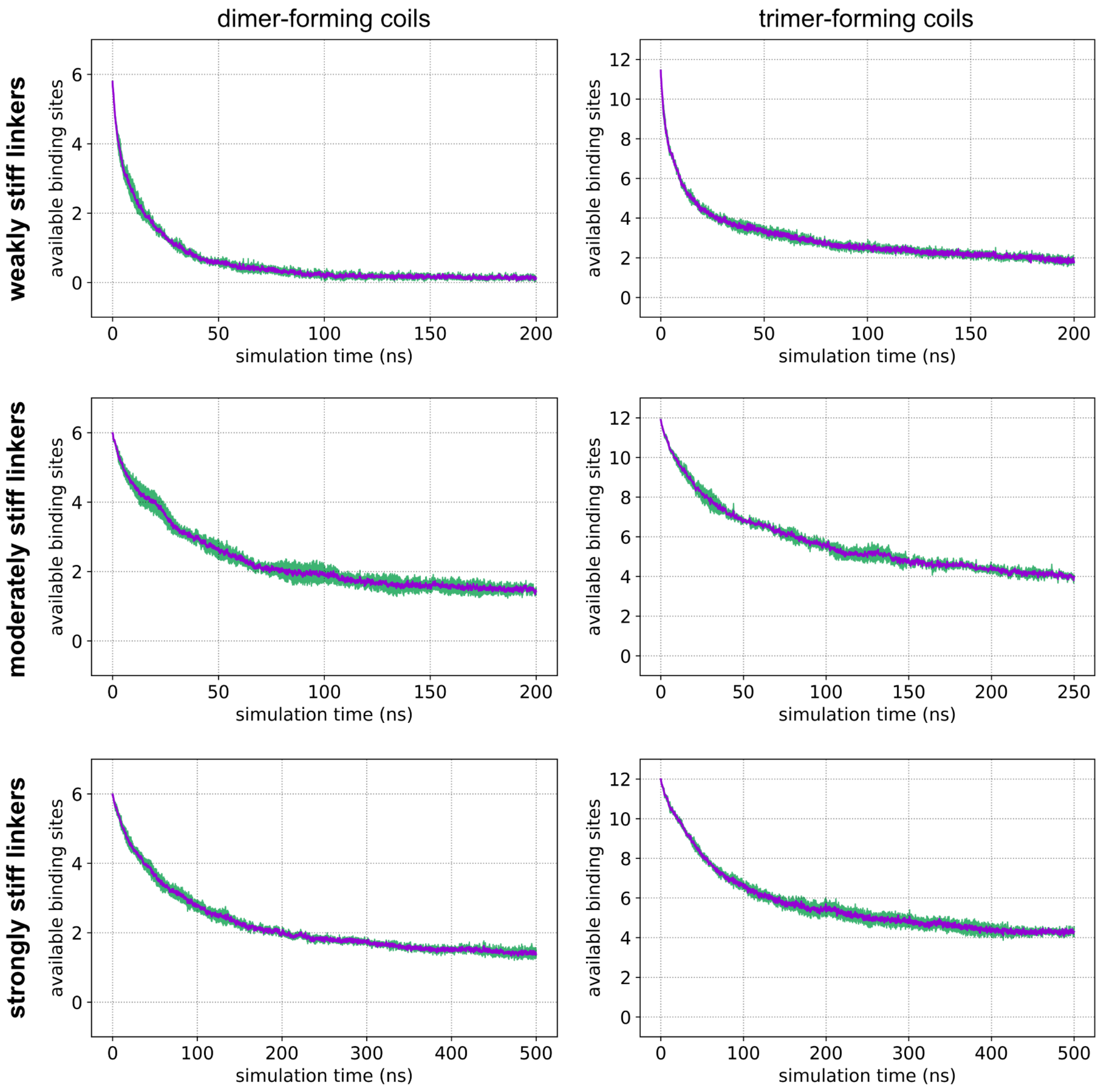

Figure S8: Number of available binding sites of single protein molecules with stiff linkers during the annealing step of the protein preparation procedure. Data are from single molecule simulations at 273 K only. The number of available binding sites of individually prepared protein molecules was averaged over all the protein molecules for a slab replicate (*n* according to Table S2), and then the average was averaged again across three slab replicates to produce the solid purple line. Green shading around the mean represents the standard deviation of three slab replicate averaged data.

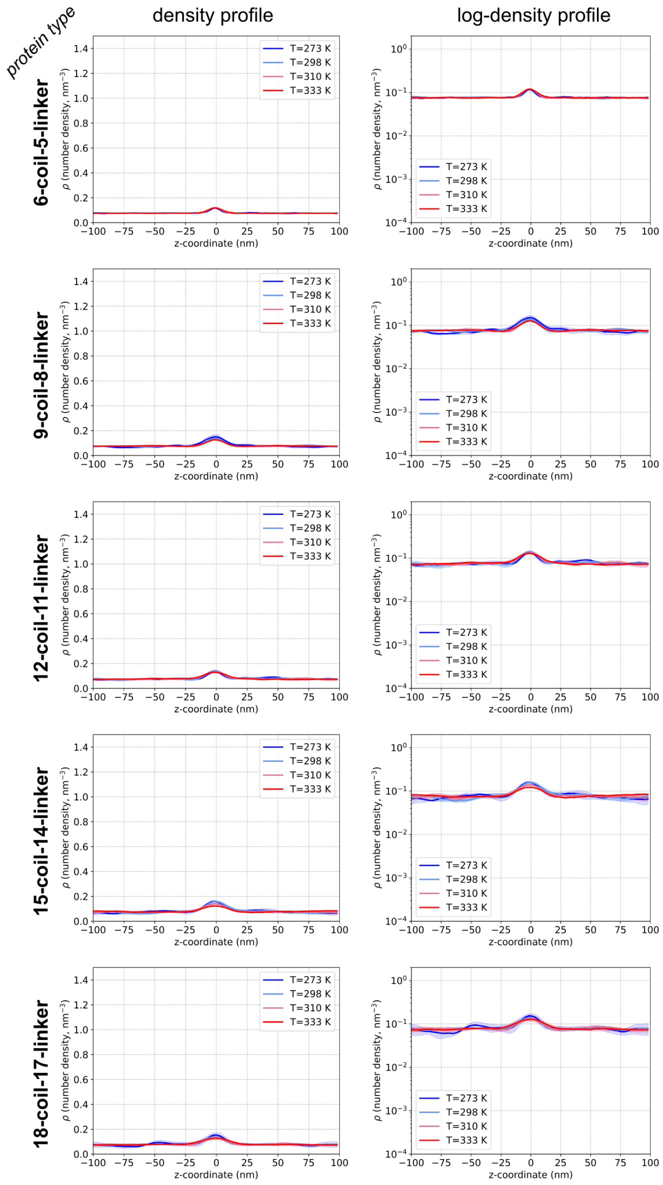

Figure S9: Density profiles from proteins with dimer-forming coils. Each row is labeled according to protein type, and columns show the density profile (left-hand side) along with the log-density transformation of the profiles (right-hand side). Solid lines represent mean, and shaded regions around the mean represent standard deviation, from three replicate simulations.

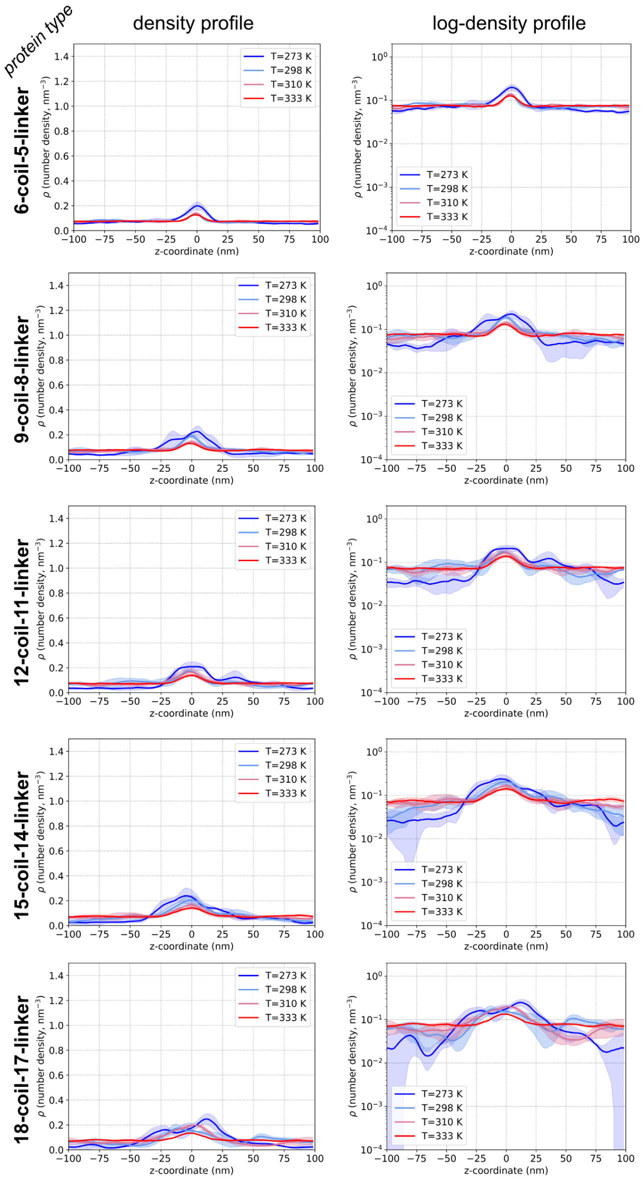

Figure S10: Density profiles from proteins with trimer-forming coils. Each row is labeled according to protein type, and columns show the density profile (left-hand side) along with the log-density transformation of the profiles (right-hand side). Solid lines represent mean, and shaded regions around the mean represent standard deviation, from three replicate simulations.

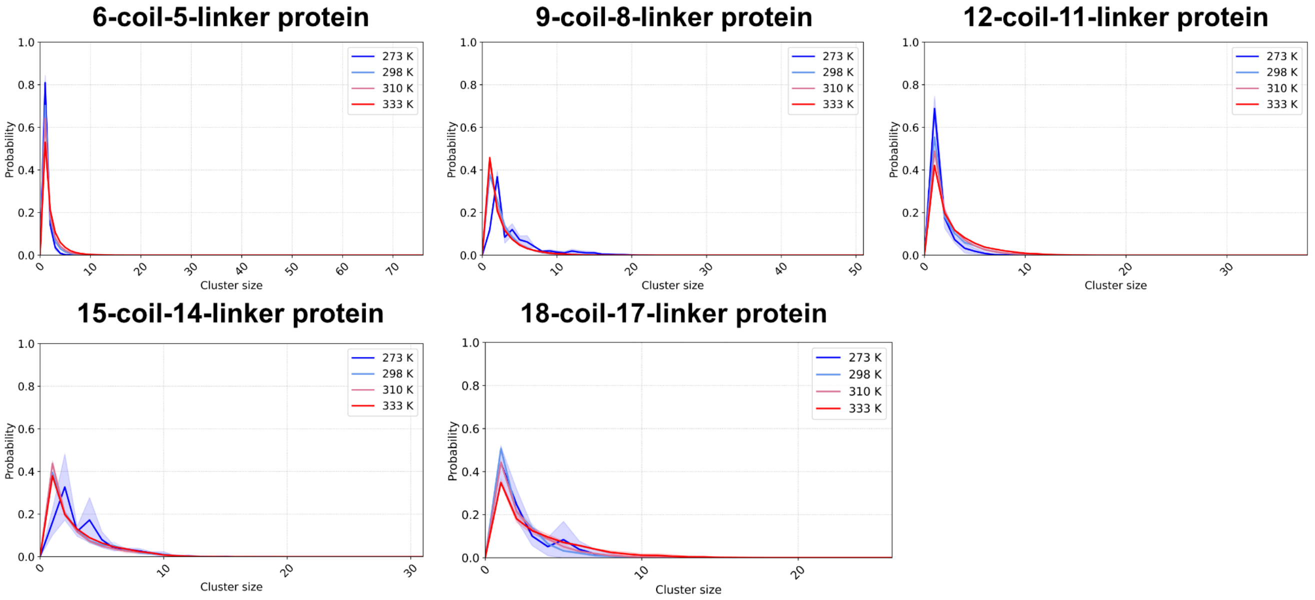

Figure S11: Molecular cluster size distributions of nonspecifically interacting dimer-forming proteins. Solid lines represent mean, and shaded regions around the mean represent standard deviation, from three replicate simulations.

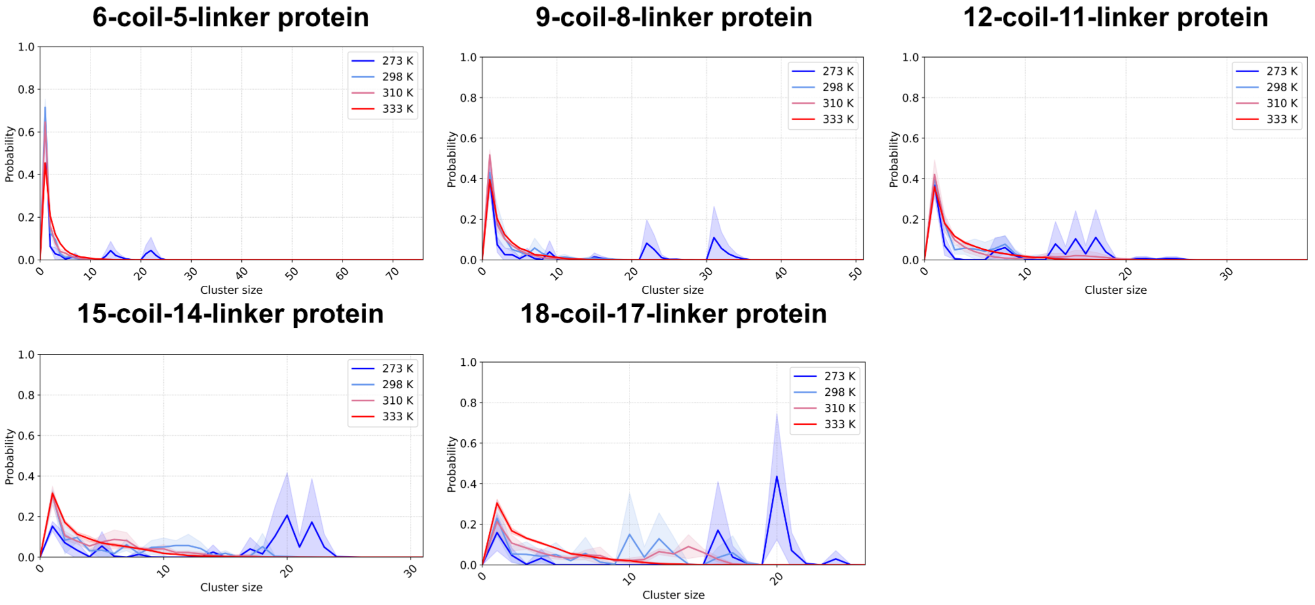

Figure S12: Molecular cluster size distributions of nonspecifically interacting trimer-forming proteins. Solid lines represent mean, and shaded regions around the mean represent standard deviation, from three replicate simulations.

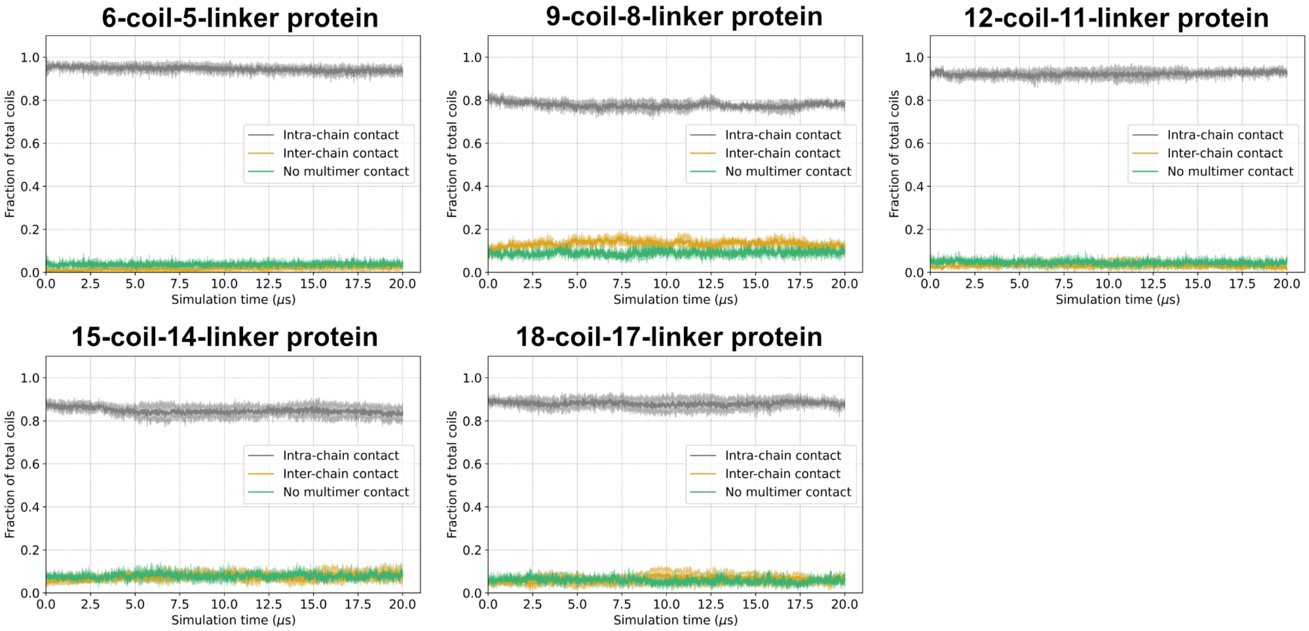

Figure S13: Types of coil interactions formed in slab simulations of dimer-forming proteins. Solid lines represent mean, and shaded regions around the mean represent standard deviation, from three replicate simulations. In some cases, e.g. "no multimer contact" classifications, the standard deviation is smaller than can be resolved on the plot.

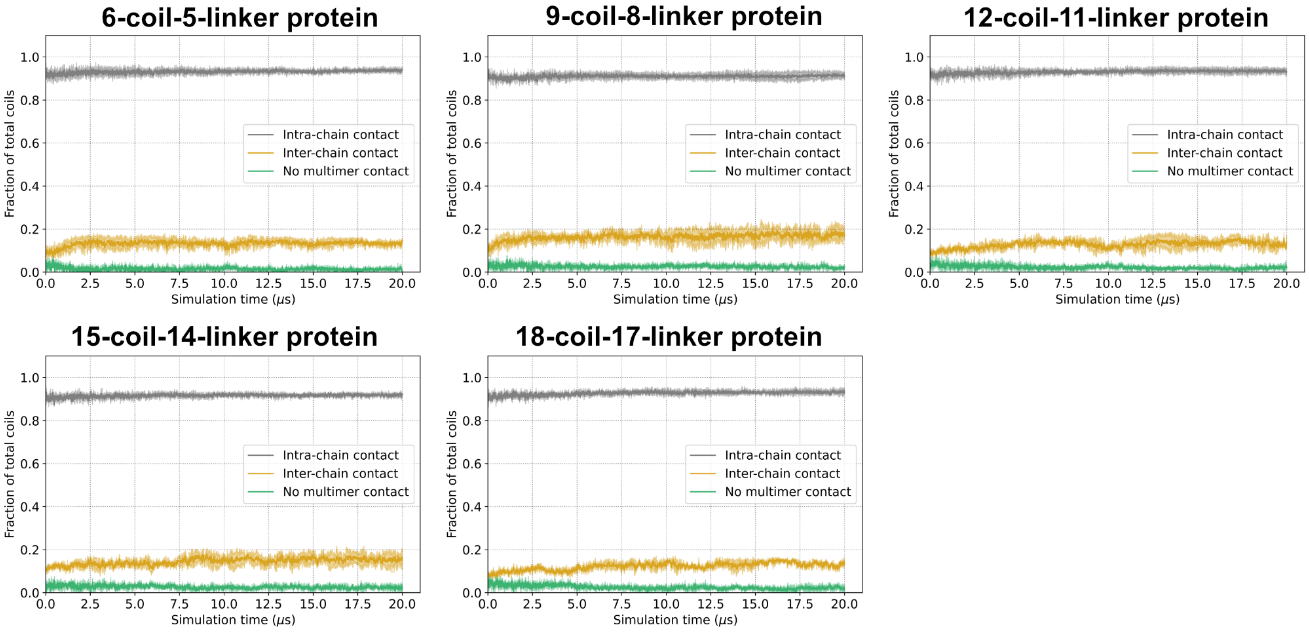

Figure S14: Types of coil interactions formed in slab simulations of trimer-forming proteins. Solid lines represent mean, and shaded regions around the mean represent standard deviation, from three replicate simulations. In some cases, e.g. "no multimer contact" classifications, the standard deviation is smaller than can be resolved on the plot.

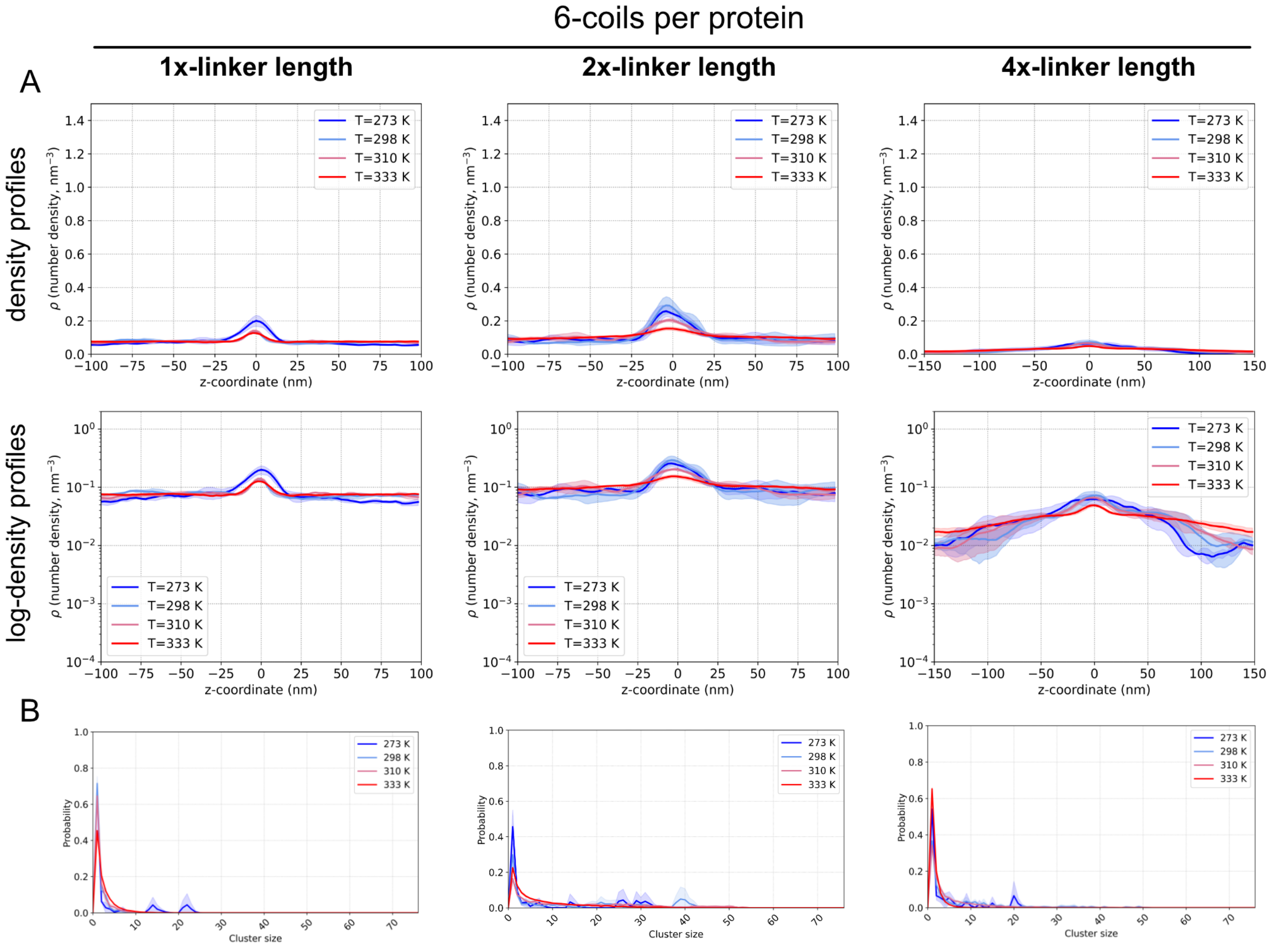

Figure S15: Density profile and molecular cluster analysis for 6-coil proteins with increasing linker length. (A) Profile densities, including log-density transformed plots, and (B) molecular cluster analyses. Each column corresponds to the linker length indicated by the column header. Profile densities and molecular cluster plots for the 1x-linker protein are the same as in Figures S10 and S12, respectively. Solid lines represent mean, and shaded regions around the mean represent standard deviation, from three replicate simulations.

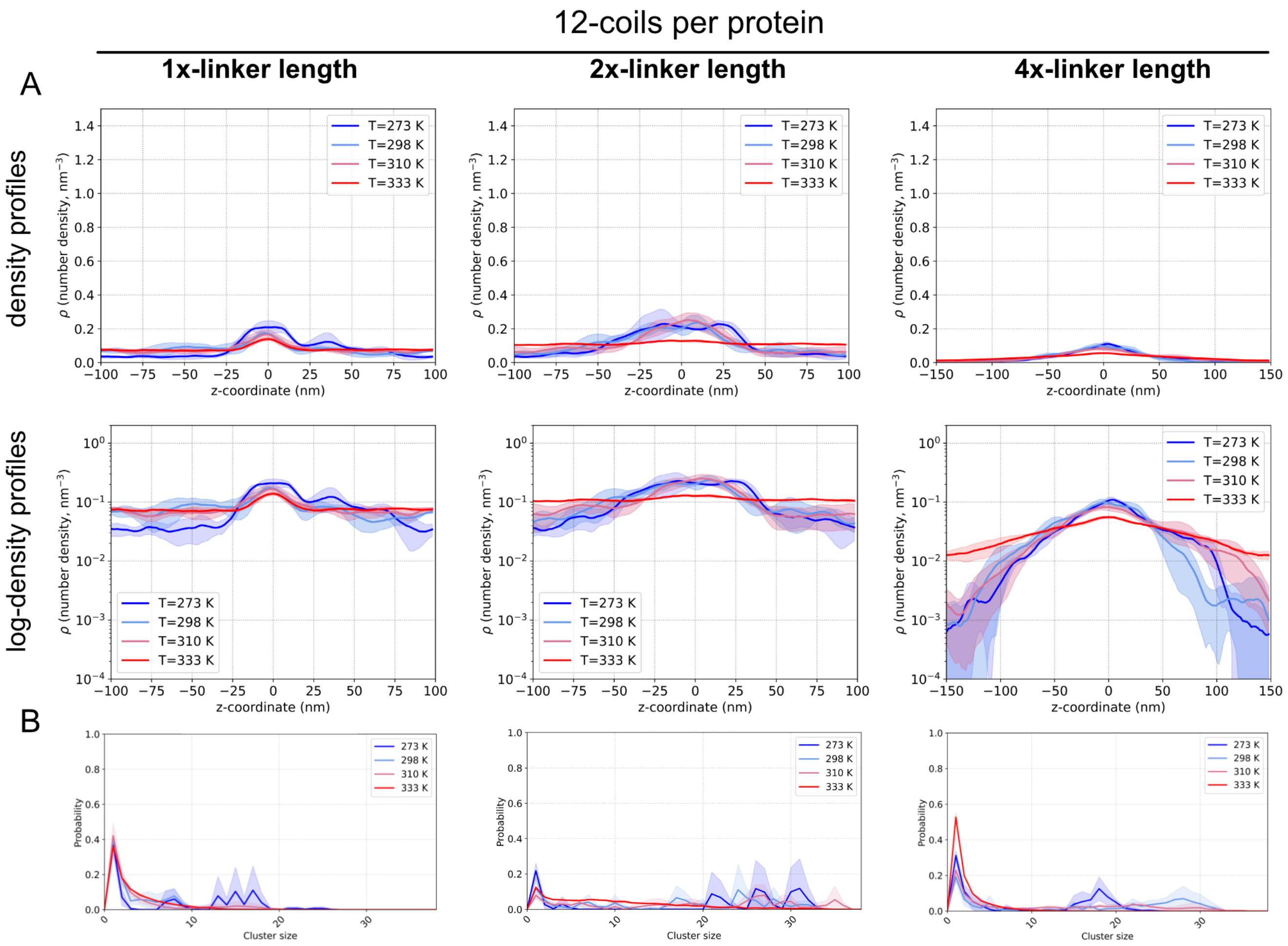

Figure S16: Density profile and molecular cluster analysis for 12-coil proteins with increasing linker length. (A) Profile densities, including log-density transformed plots, and (B) molecular cluster analyses. Each column corresponds to the linker length indicated by the column header. Profile densities and molecular cluster plots for the 1x-linker protein are the same as in Figures S10 and S12, respectively. Solid lines represent mean, and shaded regions around the mean represent standard deviation, from three replicate simulations.

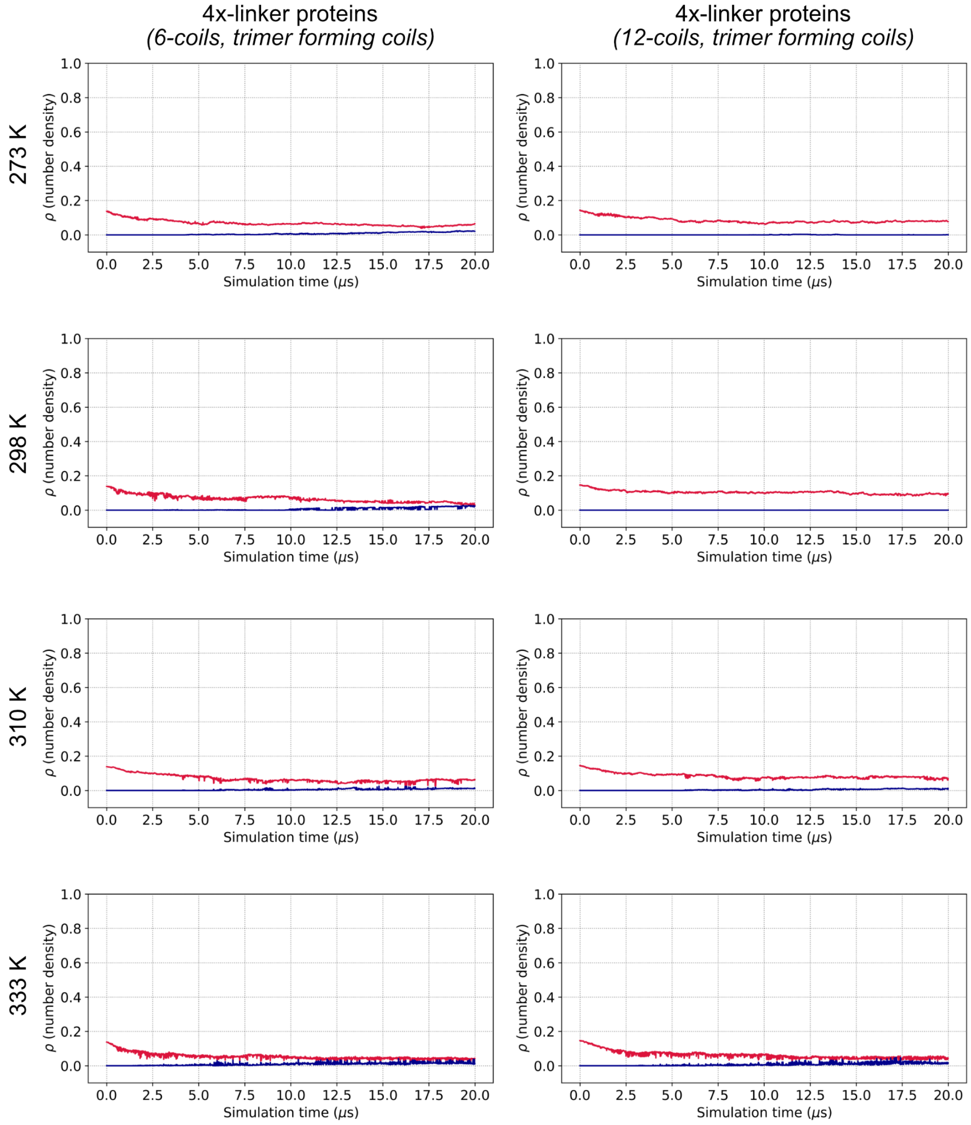

Figure S17: Proteins with 4x-long linkers reach density equilibrium within the slab simulation time. Example density equilibration plots (Methods, sec. *Determining equilibrium in slab simulations*) are shown from one set of slab replicates for the listed proteins (column titles) at each of the tested temperatures (row titles). Plots show the number density of the center of the box (red line), where a droplet would be in a phase-separated state, and the number density of the dilute regions of the box (blue line). Each plot is from one slab simulation replicate and thus no standard deviation is shown.

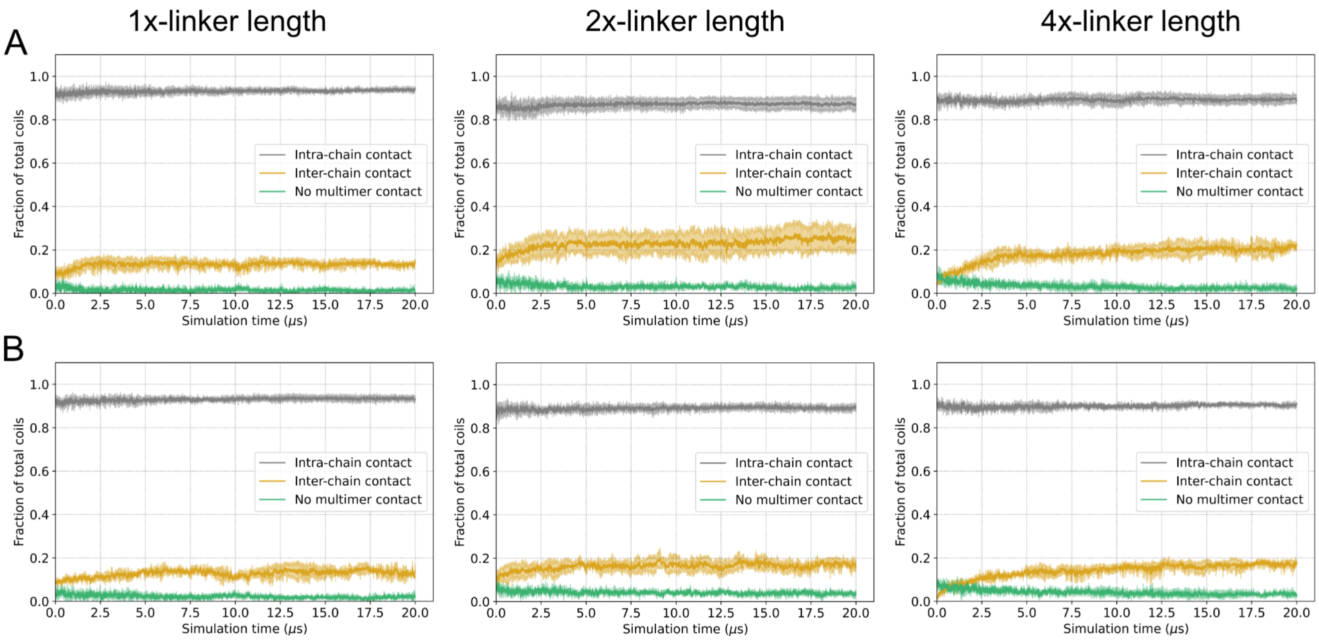

Figure S18: Types of coil interactions formed in slab simulations of proteins with increasing linker lengths. Coil contact classification plots from slab simulations of (A) 6-coil and (B) 12-coil proteins, each type of proteins with trimer-forming coils. Columns indicate the linker length as indicated by the headings. Solid lines represent mean, and shaded regions around the mean represent standard deviation, from three replicate simulations. In some cases, e.g. "no multimer contact" classifications, the standard deviation is smaller than can be resolved on the plot.

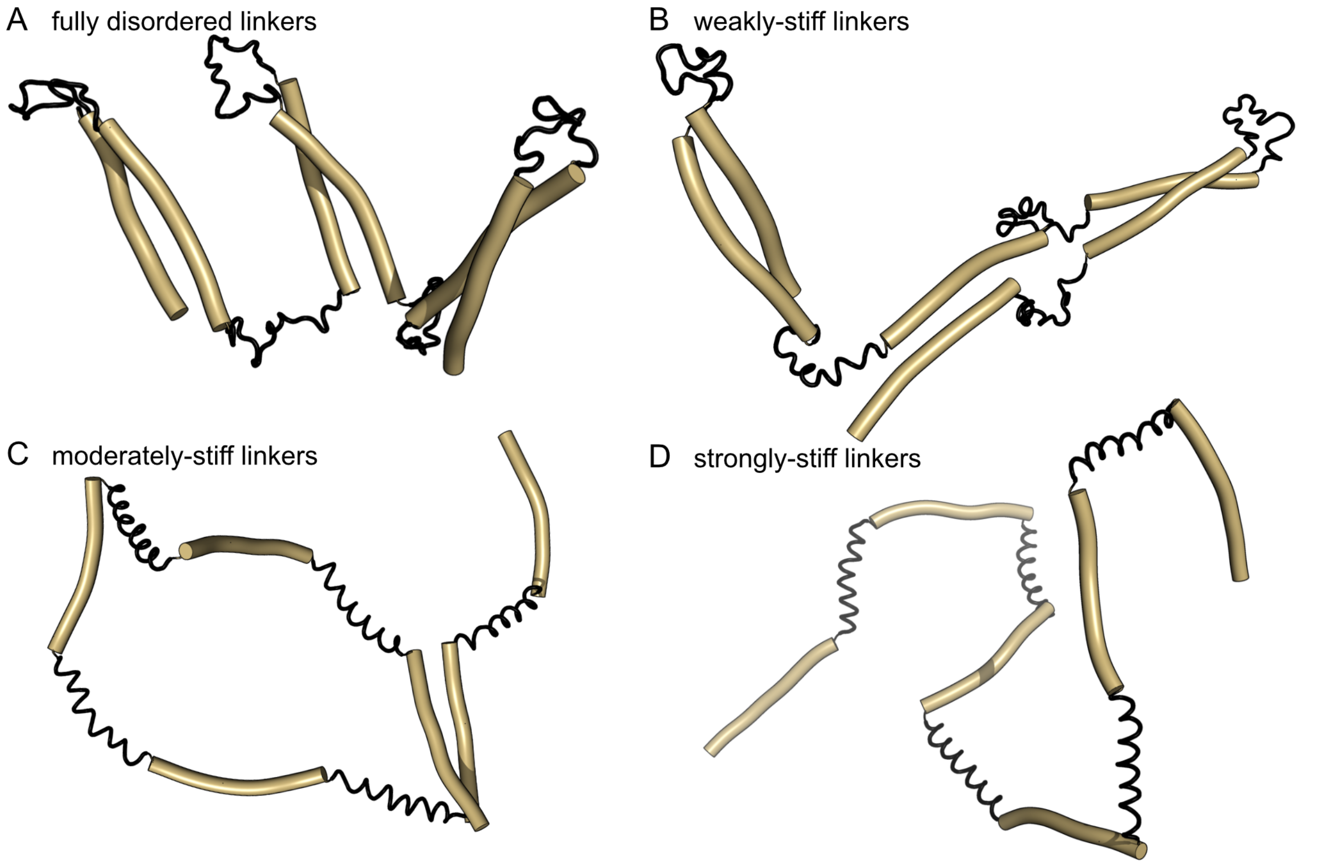

Figure S19: Increasing linker stiffness results in linkers that appear helical. Snapshots of single proteins with (A) fully disordered, (B) weakly-stiff, (C) moderately-stiff, and (D) strongly-stiff linkers to provide a physical example of linker stiffness. Snapshots of the single proteins are taken from the final configuration of slab simulations of dimer-forming coil proteins at the respective linker stiffness. Coil segments are represented as golden cylinders, and linker segments as thin black lines. Images were produced using open-source PyMOL v.2.5.0.

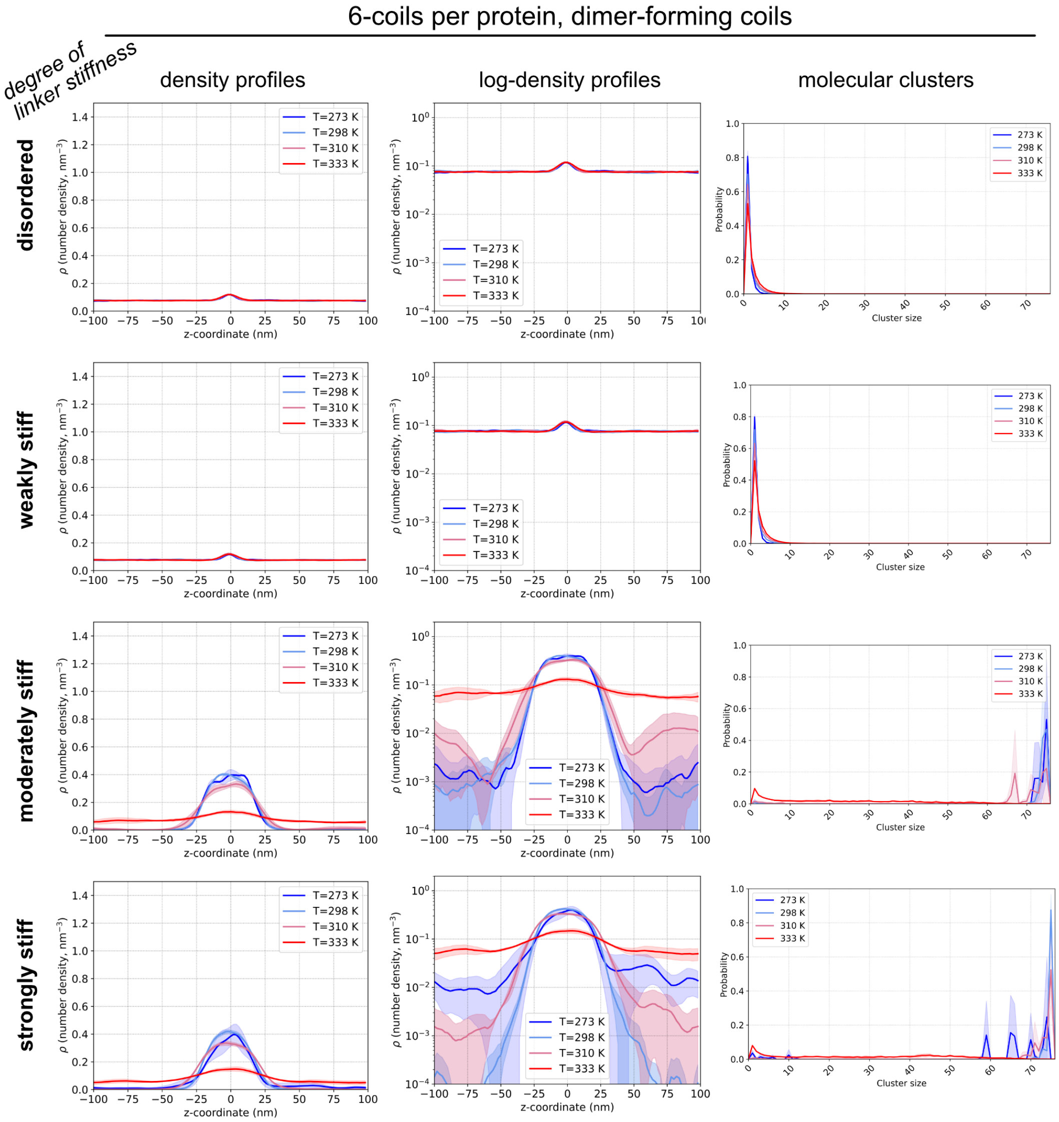

Figure S20: Density profile and molecular cluster analysis for dimer-forming 6-coil proteins with varying linker stiffness. Each row corresponds to a set of proteins distinguished by the stiffness of their linkers. The first two columns show density profiles along with a log-density transformation, and the far right column shows the molecular cluster size distributions. Solid lines represent mean, and shaded regions around the mean represent standard deviation, from three replicate simulations.

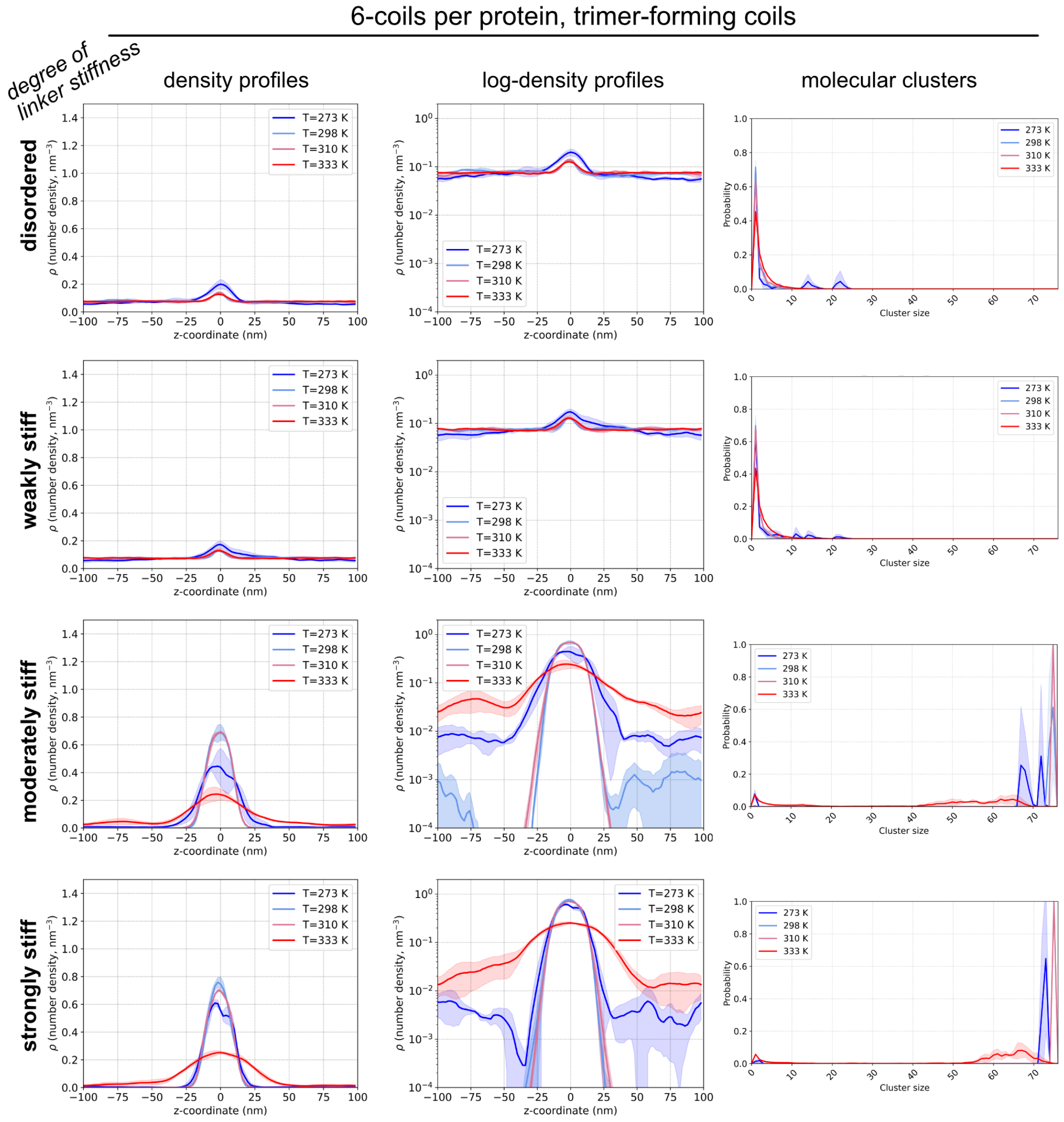

Figure S21: Density profile and molecular cluster analysis for trimer-forming 6-coil proteins with varying linker stiffness. Each row corresponds to a set of proteins distinguished by the stiffness of their linkers. The first two columns show density profiles along with a log-density transformation, and the far right column shows the molecular cluster size distributions. Solid lines represent mean, and shaded regions around the mean represent standard deviation, from three replicate simulations.

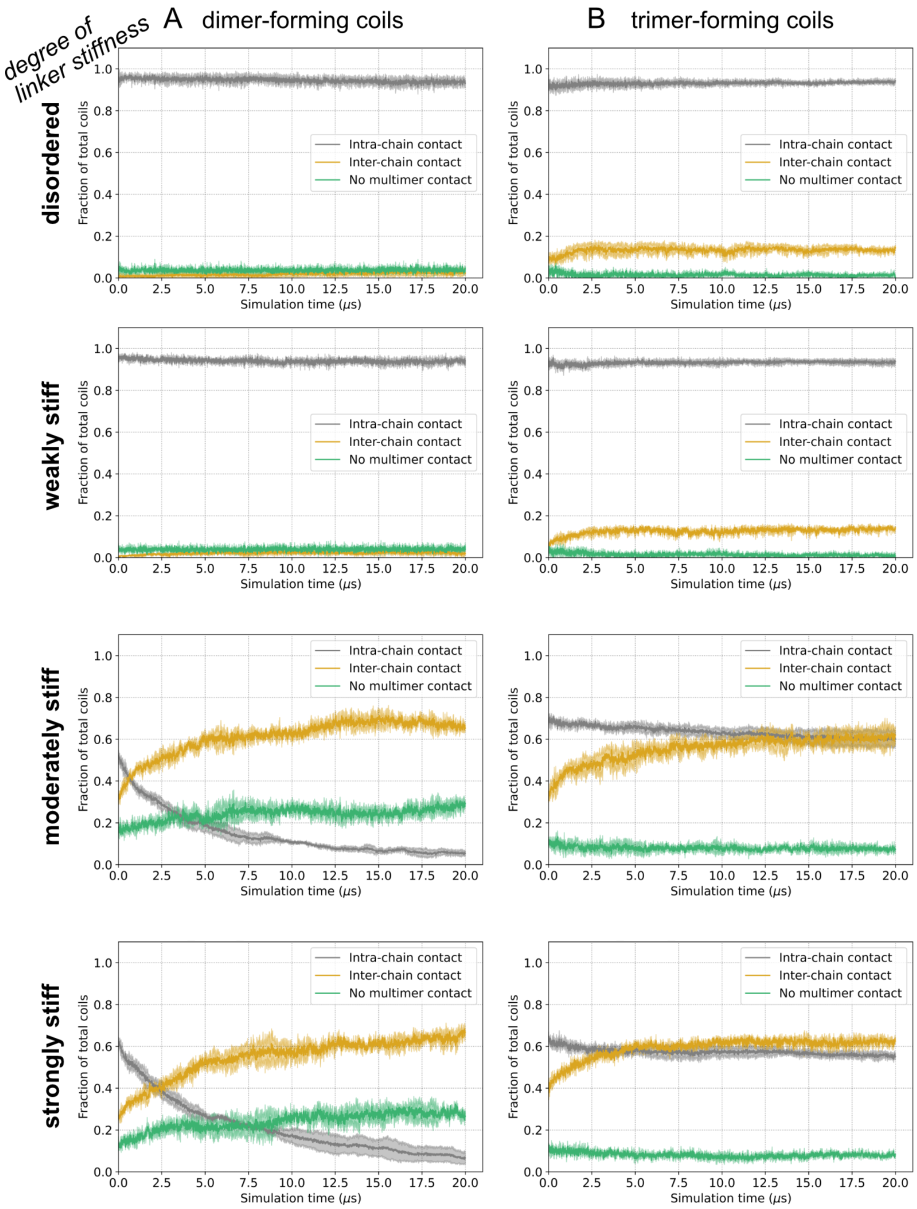

Figure S22: Types of coil interactions formed in slab simulations of dimer- and trimer-forming coil proteins with increasing linker stiffness. Coil contact classification plots from slab simulations of (A) dimer-forming and (B) trimer-forming coil proteins, with varying linker stiffness as indicated on the left-hand side separated by rows. Solid lines represent mean, and shaded regions around the mean represent standard deviation, from three replicate simulations.

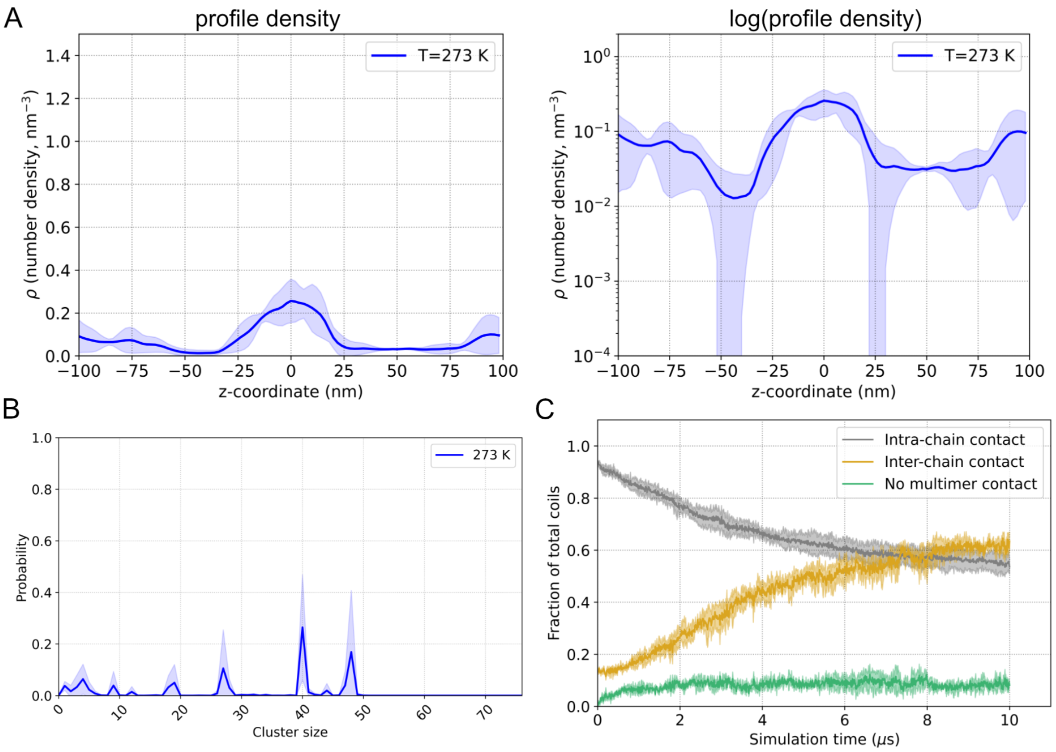

Figure S23: Swapping linker stiffness from weakly-to-moderately stiff can make protein models LLPS. (A) Density profiles (log-density transformed on right-hand side) and (B) molecular cluster size distribution analyses from an extended slab simulation of proteins that were originally with weakly-stiff linkers, but then swapped with moderately-stiff linkers. (C) Plot of coil interactions formed during the slab simulation. In all plots, solid lines represent mean, and shaded regions around the mean represent standard deviation, from three replicate simulations.

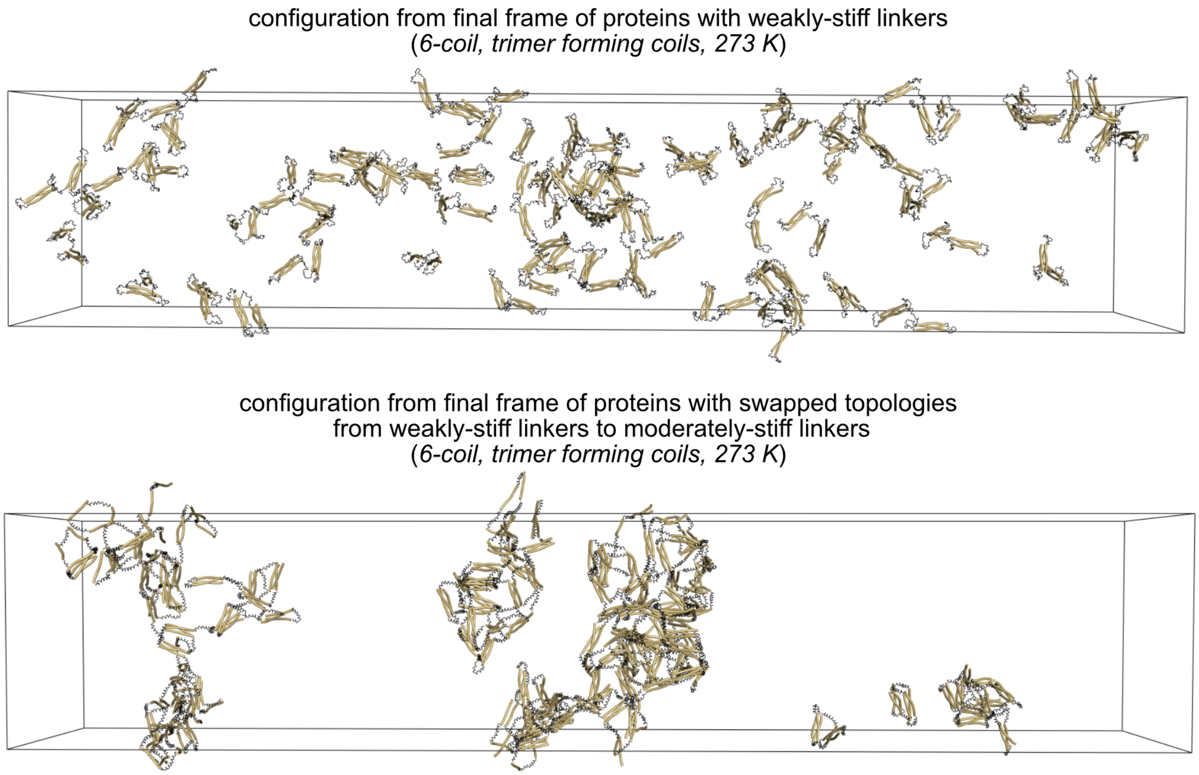

Figure S24: Proteins with topologies swapped from weakly- to moderately-stiff spontaneously form a LLPS-like cluster. The top image corresponds to the final frame configuration of a slab simulation of 6-coil timer-forming proteins with weakly-stiff linkers, used as the input for the linker-swap simulations. The bottom image corresponds to the final frame of the linker-swap simulations were proteins with originally weakly-stiff linkers used the topology of moderately-stiff linkers instead. Simulation images represent examples from a set of slab replicates. Proteins are made whole for visualization, but would wrap through the box boundaries during actual simulation. Visualizations produced using open-source PyMOL v.2.5.0.

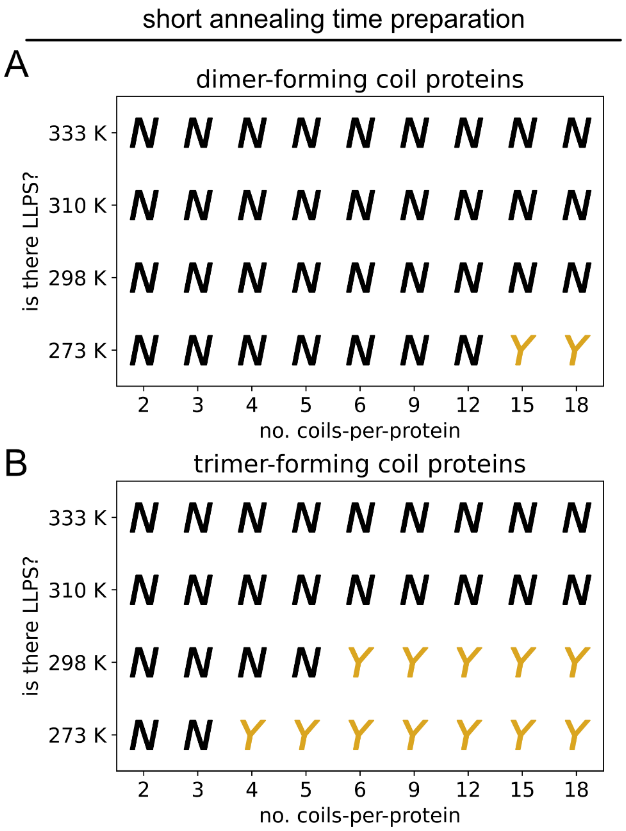

Figure S25: Proteins generated from short-annealing times have increased LLPS propensity. Phase diagrams for (A) dimer-forming, and (B) trimer-forming coil proteins with various coils-per-protein that have preferentially few intra-chain coil contacts generated from shorter preparative annealing times. Assessments of LLPS propensity were made from three replicate slab simulations.

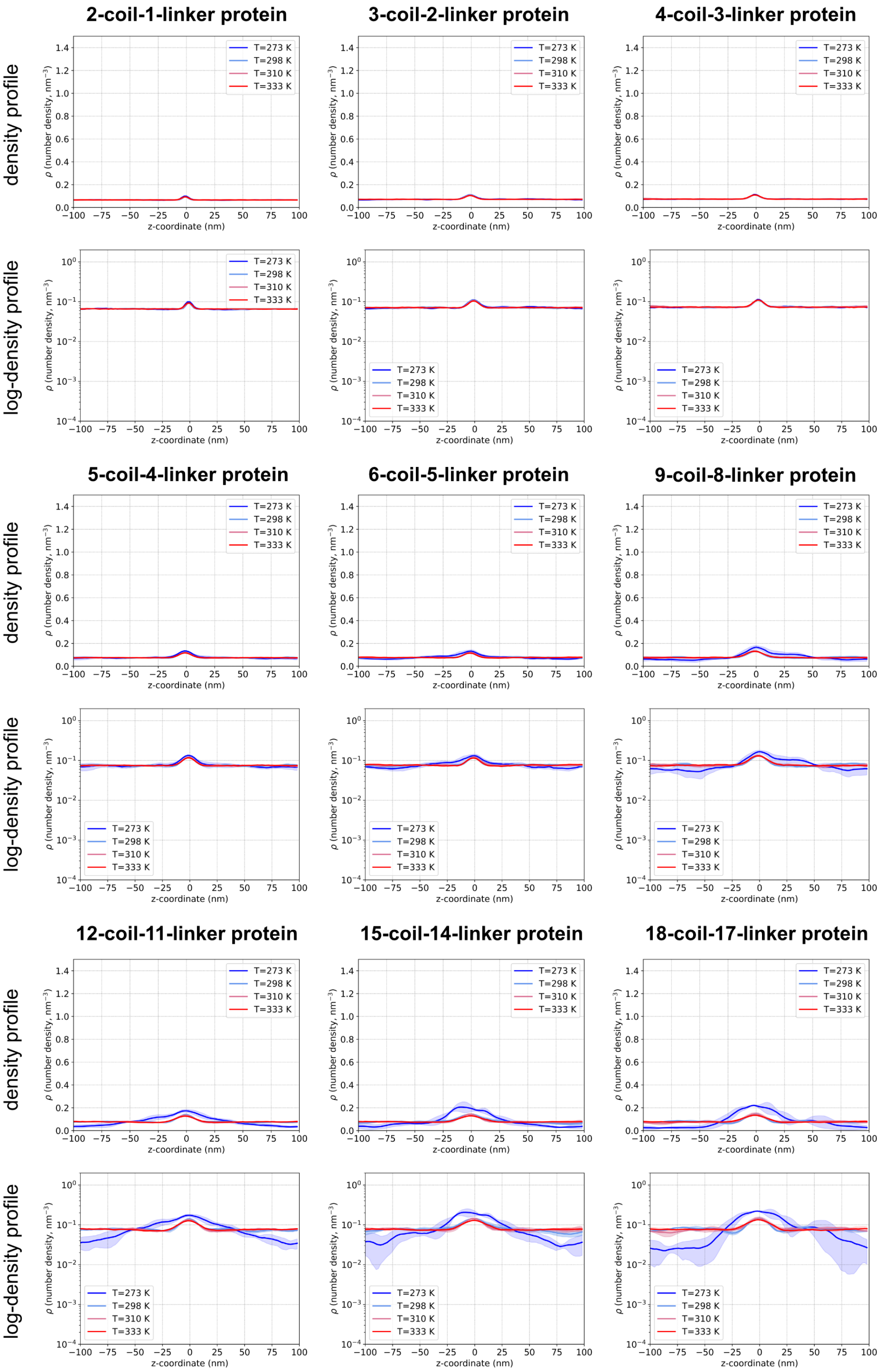

Figure S26: Density profiles from proteins with dimer-forming coils that were prepared with short annealing times. Density profile plots, along with log-density transformed plots, are shown for the studied proteins (column headers, bolded text). Solid lines represent mean, and shaded regions around the mean represent standard deviation, from three replicate simulations.

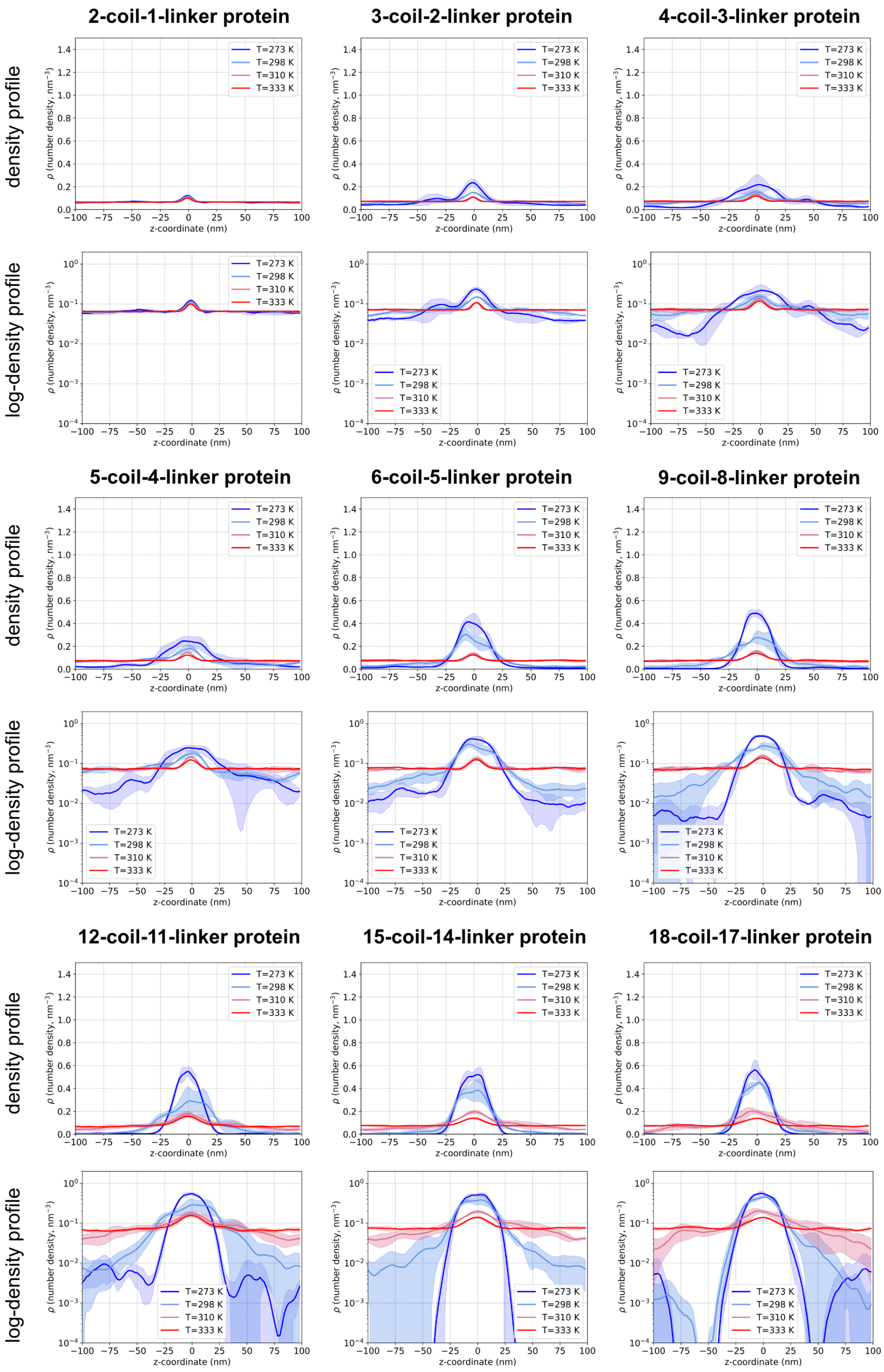

Figure S27: Density profiles from proteins with trimer-forming coils that were prepared with short annealing times. Density profile plots, along with log-density transformed plots, are shown for the studied proteins (column headers, bolded text). Solid lines represent mean, and shaded regions around the mean represent standard deviation, from three replicate simulations.

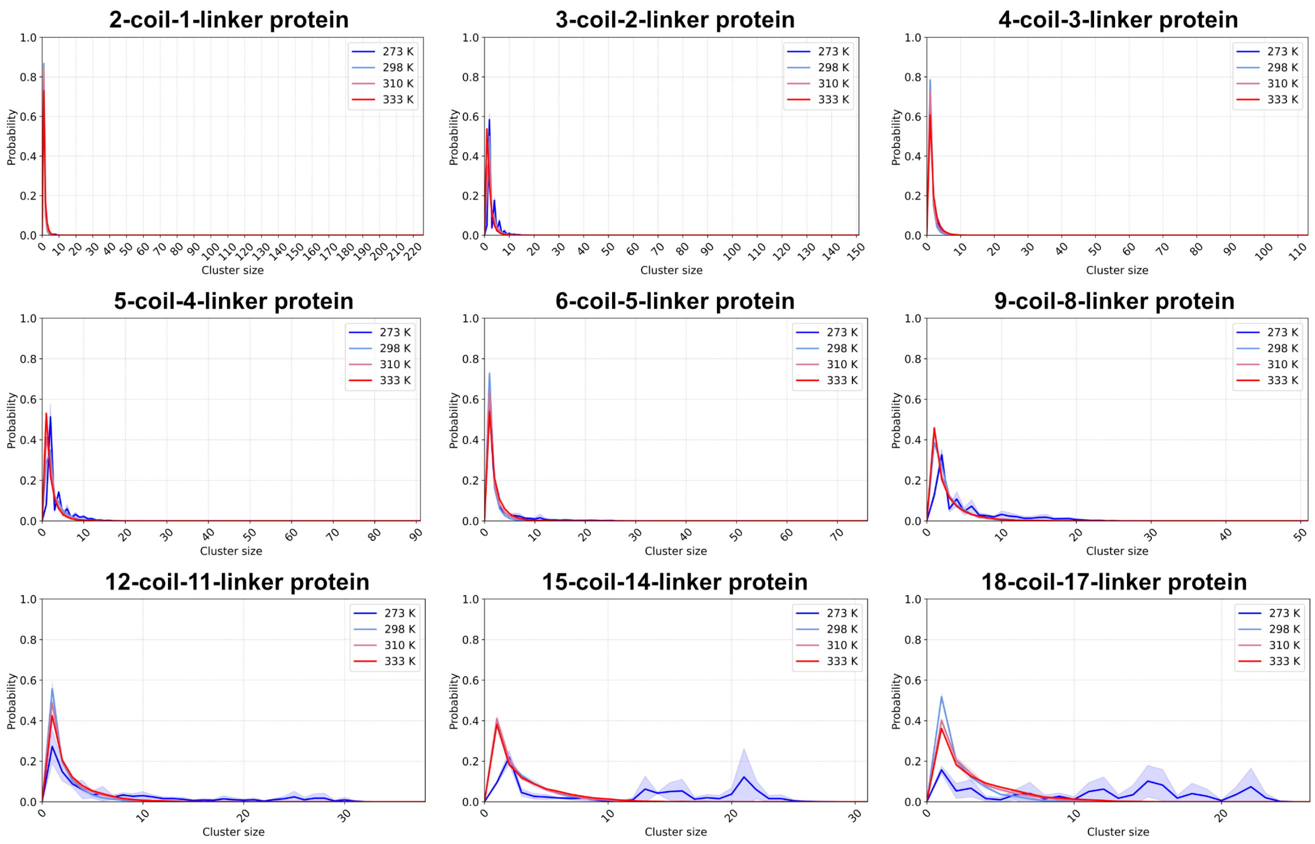

Figure S28: Molecular cluster size distributions of nonspecifically interacting dimer-forming proteins from preparations with short annealing times. Solid lines represent mean, and shaded regions around the mean represent standard deviation, from three replicate simulations.

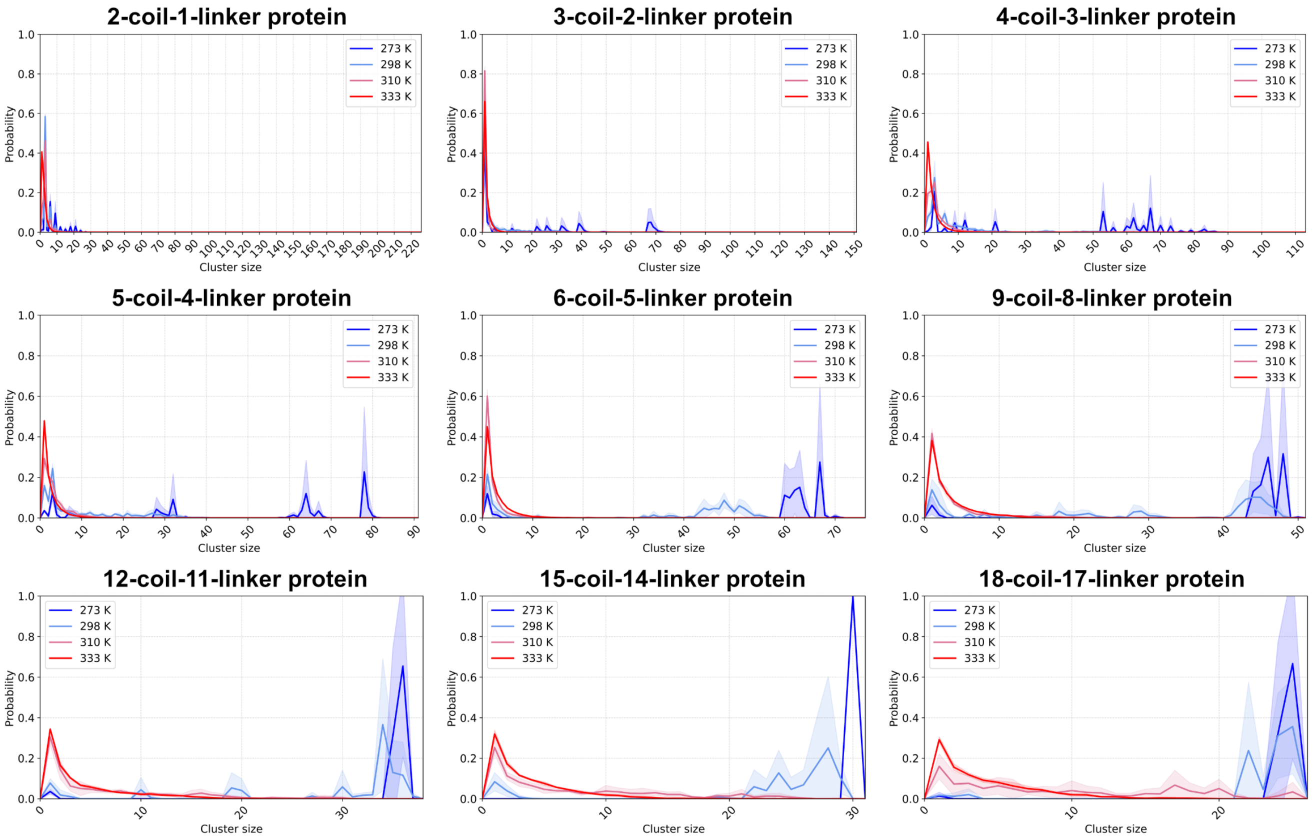

Figure S29: Molecular cluster size distributions of nonspecifically interacting trimer-forming proteins from preparations with short annealing times. Solid lines represent mean, and shaded regions around the mean represent standard deviation, from three replicate simulations.

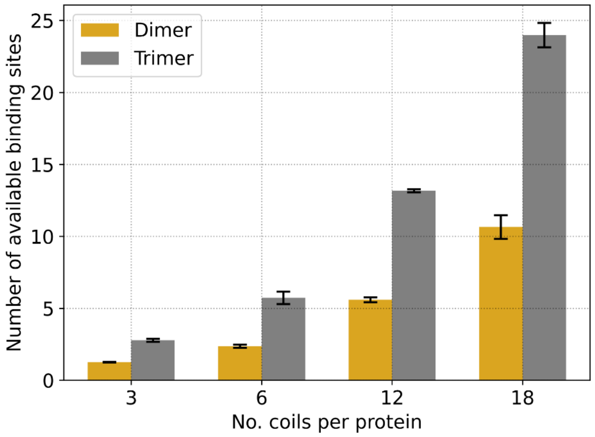

Figure S30: Trimer-forming coil proteins from short annealing preparations have a high average number of available sites for inter-chain contacts to form. Plot of number of available binding sites from protein configurations at 273 K for proteins prepared through short annealing simulations. Solid bars represent the mean, and black error bars represent the standard deviation, from replicate averaged number of available binding sites of configurations averaged over three slab simulations.

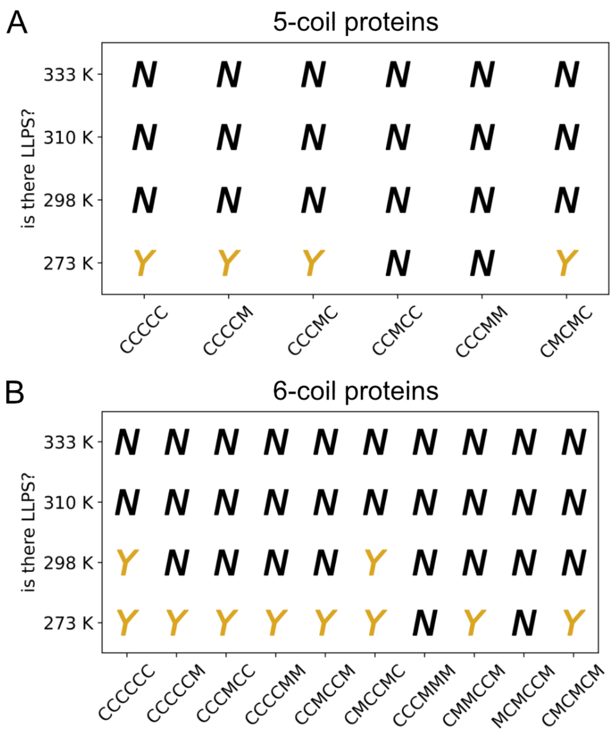

Figure S31: Coil proteins that contain mismatched coil lengths do not LLPS better than coil proteins with coils all of the same length. Phase diagrams of (A) 5-coils per protein, and (B) 6-coils per proteins, all with trimer-forming coils, with proteins that have coil segments of shorter-than-normal length. Labels on $x$-axes demonstrate the organization of coil segments in each protein, with “C" representing the default 5-heptad coils, and “M" representing the mismatched and shorter 3-heptad coils. Assessments of LLPS propensity were made from three replicate slab simulations. LLPS assessments of the no mismatch ‘CCCCC’ and ‘CCCCCC’ proteins are taken from the 5-coil and 6-coil data Figure S25B, respectively.

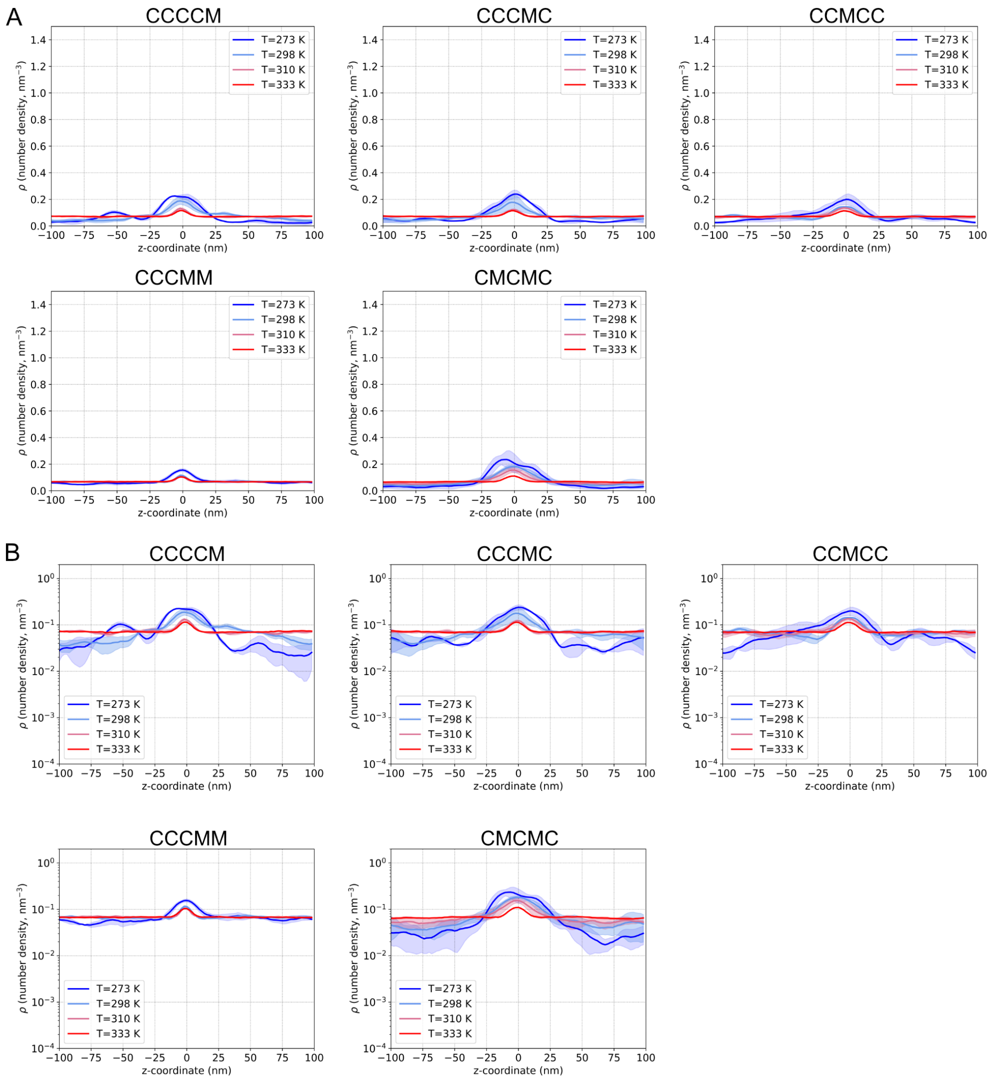

Figure S32: Density profiles from 5-coil proteins with coils of mismatched heptad length. (A) Density profile plots, along with (B) log-density transformed plots, are shown. The protein names, which are combinations of C (5-heptad coil length) and M (mismatched coil length, 3-heptads) correspond to proteins in Figure S31A. Solid lines represent mean, and shaded regions around the mean represent standard deviation, from three replicate simulations

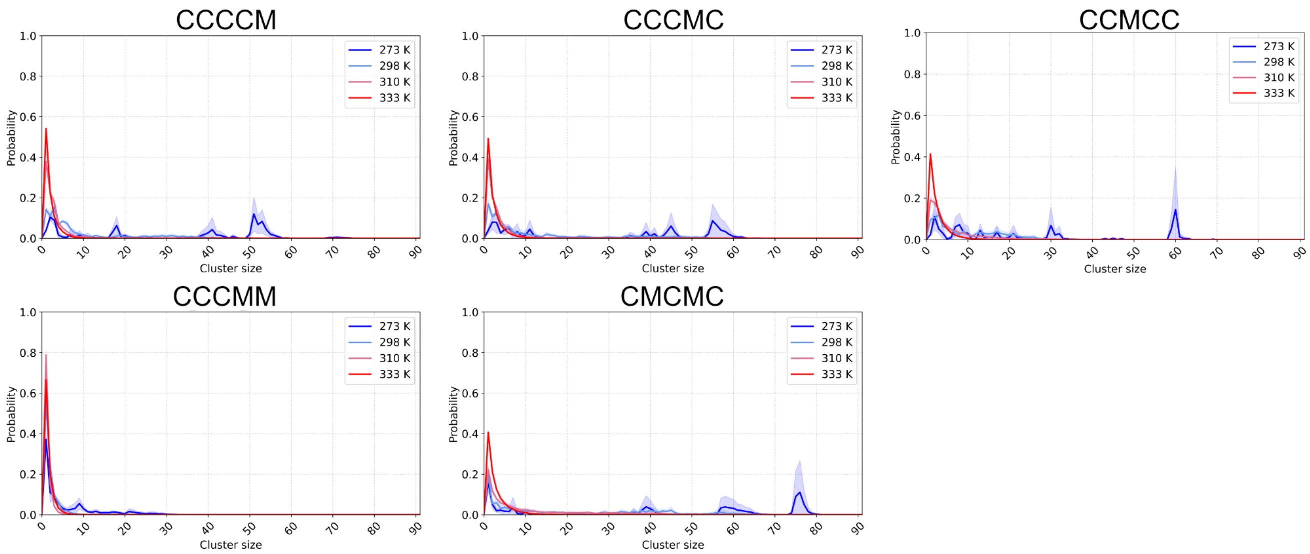

Figure S33: Molecular cluster size distributions of 5-coil proteins with coils of mismatched heptad length. The protein names, which are combinations of C (5-heptad coil length) and M (mismatched coil length, 3-heptads) correspond to proteins in Figure S31A. Solid lines represent mean, and shaded regions around the mean represent standard deviation, from three replicate simulations.

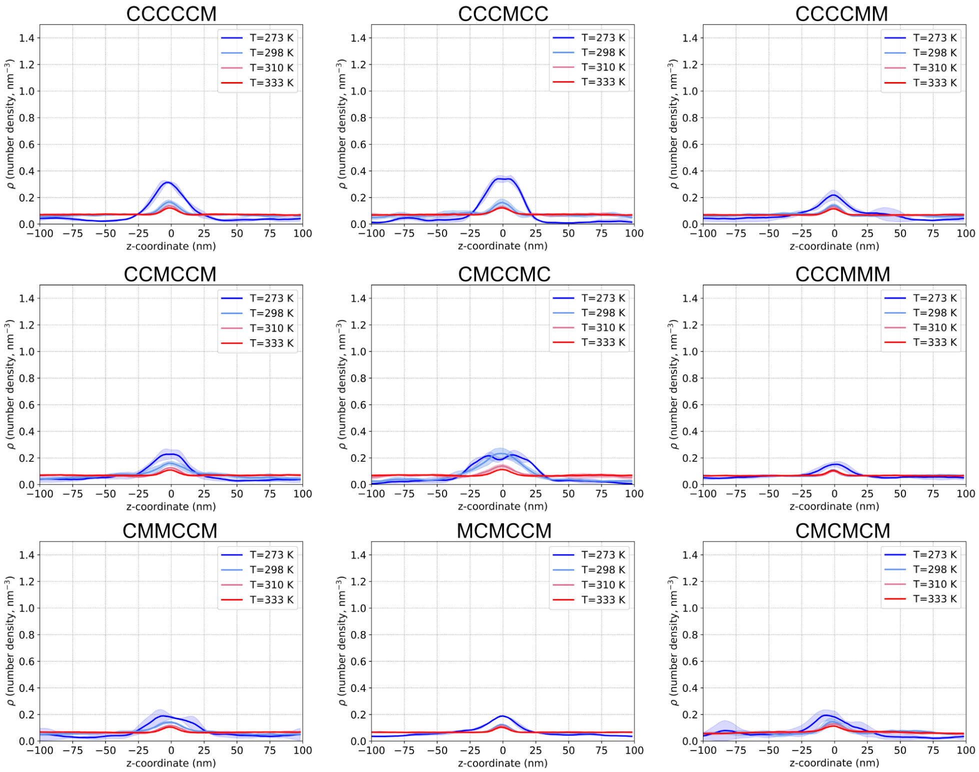

Figure S34: Density profiles from 6-coil proteins with coils of mismatched heptad length. The protein names, which are combinations of C (5-heptad coil length) and M (mismatched coil length, 3-heptads) correspond to proteins in Figure S31B. Solid lines represent mean, and shaded regions around the mean represent standard deviation, from three replicate simulations

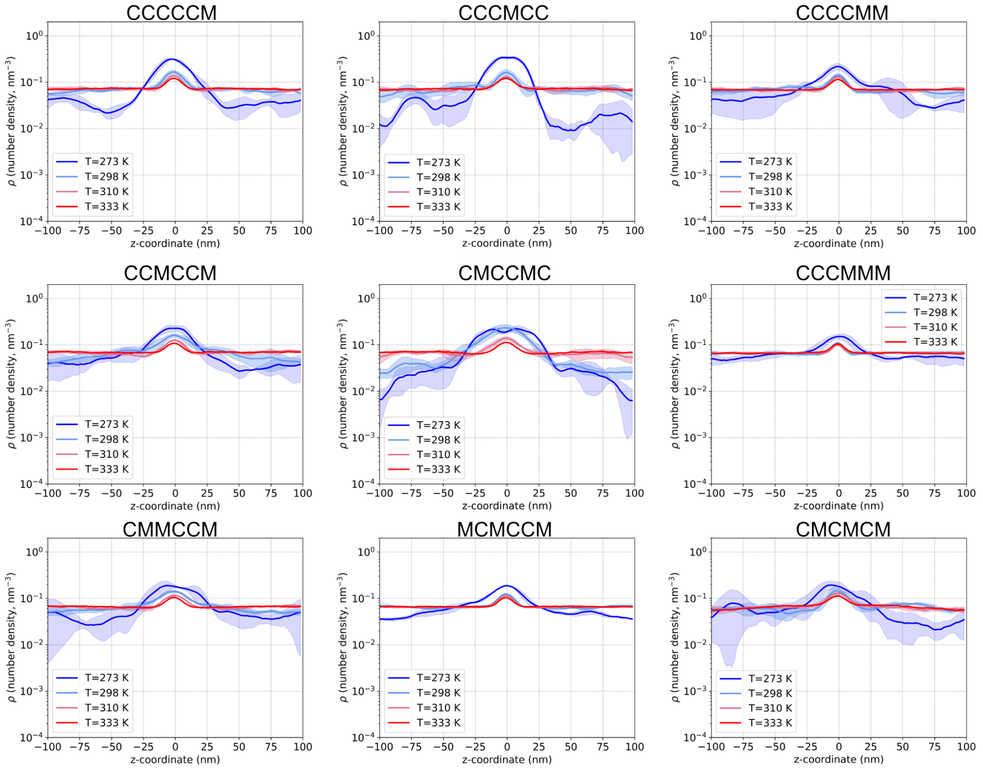

Figure S35: Log-density profiles from 6-coil proteins with coils of mismatched heptad length. The protein names, which are combinations of C (5-heptad coil length) and M (mismatched coil length, 3-heptads) correspond to proteins in Figure S31B. Solid lines represent mean, and shaded regions around the mean represent standard deviation, from three replicate simulations

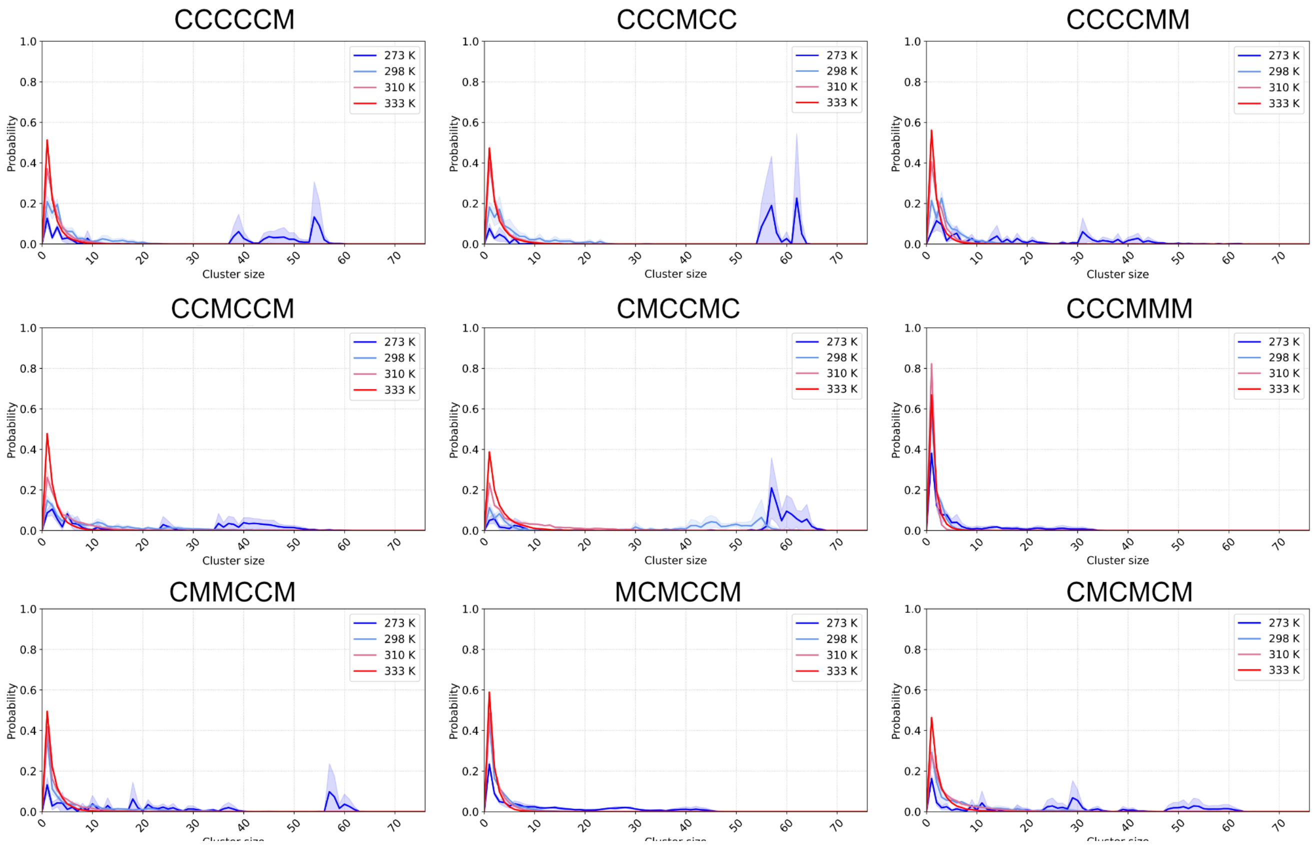

Figure S36: Molecular cluster size distributions of 6-coil proteins with coils of mismatched heptad length. The protein names, which are combinations of C (5-heptad coil length) and M (mismatched coil length, 3-heptads) correspond to proteins in Figure S31B. Solid lines represent mean, and shaded regions around the mean represent standard deviation, from three replicate simulations.

### SUPPLEMENTAL TABLES

| **Bead pairs** | **Bead class** | **$\epsilon$ (kJ/mol)** | **$\sigma$ (nm)** | **$r_{min}$ (nm)** | $C_6$ (attractive) | $C_{12}$ (repulsive) |
|---|---|---|---|---|---|---|
| **Beads specific for dimer-forming coils** | | | | | | |
| $A_{dim}$–$A_{dim}$ | coil-coil sticky | 5.5 | 0.570 | 0.640 | 7.559142E-01 | 2.597302E-02 |
| Di–Di | multimer-driving | 3.0 | 0.891 | 1.00 | 6.00 | 3.00 |
| **Beads specific for trimer-forming coils** | | | | | | |
| $A_{tri}$–$A_{tri}$ | coil-coil sticky | 4.35 | 0.570 | 0.640 | 5.978594E-01 | 2.054229E-02 |
| Tri–Tri | multimer-driving | 2.0 | 0.802 | 0.90 | 2.125764 | 5.648591E-01 |
| **Other bead interactions** | | | | | | |
| B–B | repulsive-only | 2.00 | 0.453 | 0.508 | 0.00 | 5.974044E-04 |
| B–$A_M$ | repulsive-only | 2.00 | 0.453 | 0.508 | 0.00 | 5.974044E-04 |
| B–*Multi* | repulsive-only | 2.00 | 0.453 | 0.508 | 0.00 | 5.974044E-04 |

Table S1: Nonspecific interaction parameters between the bead types used in the CC-LLPS framework. *Coil-coil sticky beads* are denoted as $A_{dim}$ for dimer- and $A_{tri}$ for trimer-specific sticky beads. $A_M$ refers generally to any of the two types of coil-coil sticky beads. *Multimer-driving beads* are denoted either Di or Tri for dimer- and trimer-specific multimer beads, respectively. *Multi* refers generally to any of the two types of multimer-driving beads. *repulsive-only beads* are denoted by the letter B. The parameters $\epsilon$ and $\sigma$ are used in the Lennard-Jones equation described in Supporting Material sec. S.II, equations 5 and 6, to generate the $C_6$ and $C_{12}$ values shown. The value $r_{min}$ is the distance between two beads at the minimum the Lennard-Jones functional and is used to calculate the corresponding $\sigma$ parameter as follows: $r_{min} = \sigma \times 2^{1/6}$. This table is reproduced and modified from Ramirez et al. (1).

| Coil segments per protein | Number of protein copies in slab simulation | Total number of coil segments |
|---|---|---|
| 2 | 225 | 450 |
| 3 | 150 | 450 |
| **4** | **112** | **448** |
| 5 | 90 | 450 |
| 6 | 75 | 450 |
| 9 | 50 | 450 |
| **12** | **37** | **444** |
| 15 | 30 | 450 |
| 18 | 25 | 450 |

Table S2: The number of protein copies, and total number of coil segments, in slab simulations for proteins with various polymeric multivalency (number of coil segments). The rows in boldface are to emphasize which slab simulations are different from the others.

| | Simulation length (ns) | |
| --- | --- | --- |
| | Hot step | Annealing step |
| Disordered linkers | 100 | 250 |
| Long linkers | | |
| *2x-long, 6-coils* | 100 | 200 |
| *2x-long, 12-coils* | 100 | 400 |
| *4x-long, 6-coils* | 300 | 400 |
| *4x-long, 12-coils* | 700 | 800 |
| Stiff linkers | | |
| *weakly stiff* | 100 | 200 |
| *moderately stiff, dimer* | 100 | 200 |
| *moderately stiff, trimer* | 100 | 250 |
| *strongly stiff* | 100 | 500 |
| Disordered linkers, short annealing time | 100 | 10 |
| Mismatched proteins | 100 | 10 |

Table S3: Single molecule simulation times for the different proteins in this study. The table is organized based on the order of appearance of the different proteins in the main text results.

Single molecule simulations

| Number of coils per protein | *type of linker* | VBT hot step | Time step (fs) hot step | VBT annealing step | Time step (fs) annealing step |
|---|---|---|---|---|---|
| 6 | 4x-long linker | `1E-9` | 10 | `1E-9` | 10 |
| 12 | 4x-long linker | `1E-9` | 10 | `1E-9` | 10 |
| 6 | weakly stiff | `1E-7` | 10 | `1E-7` | 10 |
| 6 | moderately stiff | `1E-7` | 5 | `1E-7` | 10 |

Table S4: MD parameters for single molecule simulations of coil proteins that deviate from the standard set of parameters. The values of `verlet-buffer-tolerance` (VBT) are shown as they would be written in the GROMACS molecular dynamics parameter file. The formal units for VBT are (kJ/mol/ps).

Slab simulations

| Number of coils per protein | *type of linker* | VBT – NPT, NVT, Production | Time step (fs) – NPT | Time step (fs) - - NVT, Production |
|---|---|---|---|---|
| 6 | 4x-long linker | `1E-11` | 5 | 10 |
| 12 | 4x-long linker | `1E-11` | 5 | 10 |

| Number of coils per protein | *type of linker* | VBT – EM, NPT, NVT, Production | Time step (fs) – NPT, NVT, Production |
|---|---|---|---|
| 6 | weakly stiff | `1E-11` | 10 |
| 6 | moderately stiff | `1E-11` | 10 |

Table S5: MD parameters for slab simulations of coil proteins that deviate from the standard set of parameters. The values of `verlet-buffer-tolerance` (VBT) are shown as they would be written in the GROMACS molecular dynamics parameter file. The formal units for VBT are (kJ/mol/ps).
